# Deep postnatal phenotyping of a new mouse model of nonketotic hyperglycinemia

**DOI:** 10.1101/2024.03.26.586818

**Authors:** Michael A. Swanson, Hua Jiang, Nicolas Busquet, Jessica Carlsen, Connie Brindley, Tim A. Benke, Roxanne A. Van Hove, Marisa W. Friederich, Kenneth N. MacLean, Michael H. Mesches, Johan L.K. Van Hove

## Abstract

Nonketotic hyperglycinemia due to deficient glycine cleavage enzyme activity causes a severe neonatal epileptic encephalopathy. Current therapies based on mitigating glycine excess have only limited impact. An animal model with postnatal phenotyping is needed to explore new therapeutic approaches. We developed a *Gldc* p.Ala394Val mutant model and bred it to congenic status in 2 colonies on C57Bl/6J (B6) and J129X1/SvJ (J129) backgrounds. Mutant mice had reduced P-protein and enzyme activity indicating a hypomorphic mutant. Glycine levels were increased in blood and brain regions, exacerbated by dietary glycine, with higher levels in female than male J129 mice. Birth defects were more prevalent in mutant B6 than J129 mice, and hydrocephalus was more frequent in B6 (40%) compared to J129 (none). The hydrocephalus rate was increased by postnatal glycine challenge in B6 mice, more so when delivered from the first neonatal week than from the fourth. Mutant mice had reduced weight gain following weaning until the eighth postnatal week, which was exacerbated by glycine loading. The electrographic spike rate was increased in mutant mice following glycine loading, but no seizures were observed. The alpha/delta band intensity ratio was decreased in the left cortex in female J129 mice, which were less active in an open field test and explored less in a Y-maze, suggesting an encephalopathic effect. Mutant mice showed no evidence of memory dysfunction. This partial recapitulation of human symptoms and biochemistry will facilitate the evaluation of new therapeutic approaches with an early postnatal time window likely most effective.

**Take home message:** A mouse model of nonketotic hyperglycinemia is described that shows postnatal abnormalities in glycine levels, neural tube defects, body weight, electroencephalographic recordings, and in activity in young mice making it amenable for the evaluation of novel treatment interventions.

**Author contributions:** Study concept and design: JVH, MHM, NB, KNM

Animal study data: MAS, HJ, NB, MHM, JC, CB

Biochemical and genetic studies: MAS, RAVH, MWF

Statistical analysis: NB, JVH

First draft writing: JVH, NB, MHM

Critical rewriting: MAS, NB, MHM, TAB, JC, MWF, KNM, JVH

Final responsibility, guarantor, and communicating author: JVH

**Competing interest statement:** The University of Colorado (JVH, MS, KNM, HJ) has the intention to file Intellectual property protection for certain biochemical treatments of NKH. Otherwise, the authors have stated that they had no interests that might be perceived as posing a conflict or bias to this subject matter.

**Funding support:** Financial support is acknowledged form the NKH Crusaders, Brodyn’s Friends, Nora Jane Almany Foundation, the Dickens Family Foundation, the Lucas John Foundation, Les Petits Bourdons, Joseph’s Fund, the Barnett Family, Maud & Vic Foundation, Lucy’s BEElievers fund, Hope for NKH, Madi’s Mission NKH fund, and from Dr. and Ms. Shaw, and the University of Colorado Foundation NKH research fund. The study was supported by a grant (CNS-X-19-103) from the University of Colorado School of Medicine and the Colorado Clinical Translational Science Institute, which is supported by NIH/NCATS Colorado CTSA Grant Number UL1 TR002535. Contents are the authors’ sole responsibility and do not necessarily represent official NIH views. All funding sources had no role in the design or execution of the study, the interpretation of data, or the writing of the study.

**Ethics approval on Laboratory Animal Studies:** Mouse studies were carried out with approval from the Institutional Animal Care and Use Committee of the University of Colorado Anschutz Medical Campus (IACUC# 00413).

**Data sharing statement:** The data that support the findings of this study are available from the corresponding author upon reasonable request.

## 1 INTRODUCTION

Nonketotic hyperglycinemia (NKH) is a genetic infantile epileptic encephalopathy with an incidence of 1:76,000 that is caused by deficient activity of the glycine cleavage enzyme system (GCS).^1-3^ The GCS catalyzes the breakdown of glycine with tetrahydrofolate (THF) into CO_2_, NH_3_, and 5,10-methylene-THF (Figure 1A). This mitochondrial enzyme is present in liver, placenta, and brain, primarily in astrocytes. The GCS comprises 4 subunits (P, T, H, and L) (Figure 1A). In 80% of patients with NKH, deficient GCS activity is caused by pathogenic loss of function variants in the *GLDC* gene encoding the P-protein, while the remaining 20% of patients have defects in the *AMT* gene encoding the T-protein, and in very rare patients the defect resides in the *GCSH* gene encoding the H-protein.^1,4^ Most patients have the severe form of NKH and present with neonatal epileptic encephalopathy, absent psychomotor development, spasticity, and therapy-resistant epilepsy.^2,5,6^ An attenuated form of NKH is present in 15% of patients characterized by developmental progress and treatable or no epilepsy. Human neuropathology reveals persistent vacuolating myelinopathy, variable astroglial reaction, without reported changes in oligodendroglial cells or neurons.^7-14^ Brain atrophy only develops in older severely affected patients.^15-18^

**Figure 1:**
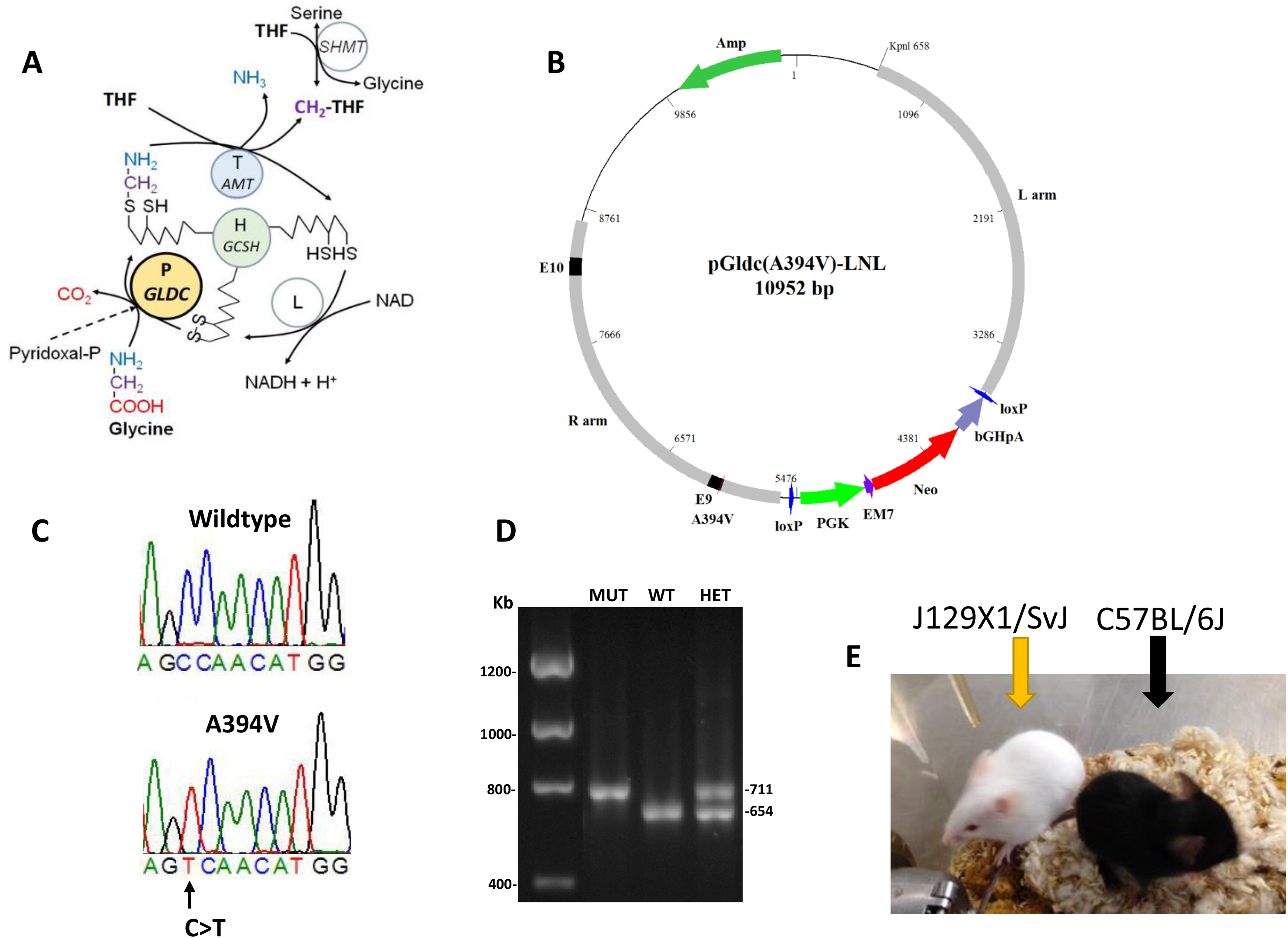
Genetics of the p.A394V mouse model. **A**. The glycine cleavage enzyme system comprises the P-protein (*GLDC* gene), the T-protein (*AMT* gene), the H-protein (*GCSH* gene), and the L-protein. **B**. The vector contains a genomic part of the mouse *Gldc* gene in a C57BL/6J background, and a neomycin and PGK selective system flanked by loxP sites. **C**. Sequencing of exon 9 shows a homozygous change of C>T in the mutant mouse. **D**. The mutant allele has a larger size than the WT allele, which was used for genotyping. **E**. Two colonies of mutant mice in the C57BL/6J strain (black coat color) and in the J129X1/SvJ strain (white coat color) were maintained in congenic status.

Deficient GCS activity results in increased circulating and brain glycine levels.^3,19^ Most past hypotheses of the pathophysiology have focused primarily on neurotoxicity caused by excess glycine,^20-22^ or on end-product deficiency of glycine derived one-carbon units to folate via 5,10-methylene-THF.^23-25^ Current therapy is exclusively focused on mitigating the effects of excess glycine and comprises glycine reduction strategies using benzoate or ketogenic diet^26-28^ and dextromethorphan to decrease putative excessive neurotransmission at the N-methyl-D-aspartate receptor.^29,30^ Early treatment moderately improves outcomes in attenuated NKH,^31^ but this treatment is ineffective in changing the outcome in severe NKH, even when initiated neonatally.^32,33^ Overall, prognosis in NKH is so dismally poor that many parents withdraw care in the neonatal period.^34^

Yet, there is reason for optimism that treatment is possible. Subjects develop increasing spasticity and worsening seizures after the first 3 months of life indicating a window of opportunity for treatment in early infancy.^2,6^ In an extensive brain imaging study,^35^ the only documented malformation was rare obstructive aqueductal hydrocephalus.^36^ The corpus callosum had decreased volume and lower growth in severely affected compared to attenuated NKH patients, indicative of impaired white matter development.^35^ Thus, imaging studies indicate a problem of brain growth rather than malformation.^35^ Effective treatment that restores brain growth may result in substantial clinical improvement. Developing new therapeutic options based on improved insights into the pathophysiology of NKH is crucial, which requires studies in mouse models.

A few mouse models of NKH have been described previously. A dominant negative transgene activated in neural stem cells results in microcephaly, early demise (<38 d), and fatal status epilepticus,^37^ whereas a mild dominant negative transgene with 29-33% residual activity exhibits some behavioral changes and longer convulsive phases after electroshock.^38,39^ A gene trap in the *AMT* gene on a C57BL/6 (B6) background with no residual activity was embryonically lethal due to neural tube defects.^40^ Unfortunately, these mouse models are no longer available.

Two gene traps in *Gldc* in intron 2 (GT1 with 5-15% normally spliced mRNA)^24^ and intron 9 (GT2 without normally spliced mRNA),^25^ on a B6 background exhibit neural tube defects, primarily exencephaly, 24% in GT1 and 57% in GT2. GT1 mice that are born alive exhibit a 55% lethality between 2 and 12 weeks of age, half from hydrocephalus and the other deaths unexplained.^24^ Biochemically, these mice have elevated glycine and decreased forms of methylated folate. Prenatal treatment with formate prevents neural tube defects and hydrocephalus and improves brain growth.^24,25,41^ Two missense *Gldc* variants were generated on the B6 mouse strain, a severe model with a complex double mutant p.Gly561Arg and p.Ser562Phe, and a milder model p.Ala394Val.^42^ In the severe double mutant, less than 10% of expected mice are born alive, whereas the p.Ala394Val mice have a 18% decreased birth rate and a 31% hydrocephalus rate, both of which are substantially improved with prenatal formate treatment. All these mice have elevated glycine levels.

In the present study, we describe a new mouse model homozygous for the missense variant *Gldc* p.Ala394Val homologous to the recurring human missense p.Ala389Val. To account for the effect of genetic background, the mouse model was made congenic on 2 backgrounds: B6 and 129X1/SvJ (J129). The mouse model phenotype was extensively characterized for glycine levels, survival, growth, brain electrophysiology, and neurobehavioral testing.

## 2 METHODS

### 2.1 Mouse Models

Mouse studies were carried out with approval from the Institutional Animal Care and Use Committee of the University of Colorado Anschutz Medical Campus (IACUC# 00413). To generate the transgenic mice, the genomic sequence around exon 9 from the B6 mouse strain was cloned into a homologous cloning vector and the *Gldc* p.Ala394Val mutation introduced by site-directed mutagenesis (Figure 1B). The vector was transfected for integration using classic techniques into E17.1 embryonic stem cells, which are derived from an F1 hybrid of C57BL/6J and J129X1/SvJ followed by removal of selection markers (see supplemental methods).

Founder mice were backcrossed in two separate colonies, B6 and J129, using a speed congenic approach in two separate colonies to evaluate the impact of strain background via asymptomatic heterozygote intercrosses to avoid negative selective pressure on the phenotype.^43,44^ The mutation was traced by polymerase chain reaction, and verified by Sanger sequencing (Figure 1C,D).

### 2.2 Biochemistry

Mutant P-protein activity measured in the glycine exchange assay was recorded as a percentage of wild-type (WT).^5,45^ Initial *in vitro* measurements were performed on COS-7 cells transfected with a plasmid containing human WT or *GLDC* p.A389V, or with mouse *Gldc* WT or *Gldc* p.A394V.^5^ Glycine cleavage enzyme activities were also measured in brain and liver tissue from the mouse model as described in the supplemental methods. Protein levels were determined using Western blot analysis of the liver, cortex, cerebellum, hippocampus, pons, and brainstem as described in the supplemental methods. Glycine levels were determined in mouse plasma and brain tissue (cortex, cerebellum, and hippocampus) using reversed phase high-pressure liquid chromatography as described in the supplemental methods.^46,47^

### 2.3 Symptoms, weight, and survival

Mice were maintained on regular chow with body weight recorded weekly for up to 90 days, and hydrocephalus or health distress was recorded. Glycine challenge of the mice was performed by the addition of 3% or 5% (w/v) glycine to drinking water provided *ad libitum* as indicated in the supplemental methods. At the end of each experiment, tissues were dissected from euthanized mice for biochemical analyses. Mice with hydrocephalus were excluded form subsequent functional studies.

### 2.4 Electrode implantations and EEG recording

Detailed methods for electrode implantation, electroencephalographic (EEG) recording and analyses are provided in the supplemental methods. In brief, subdural electrodes for EEG recording were implanted at 5 weeks of age bilaterally over the sensorimotor cortices and local field potential depth electrodes were placed bilaterally in the dorsal hippocampal area CA1 with reference and ground surface electrodes placed over the cerebellum.^48-50^ After recovery from surgery for 7 days (d), continuous video-EEG recordings were obtained for 14 d with mice receiving regular food and water *ad libitum*, and then for an additional 7 d with glycine (5% w/v) added to the drinking water. EEG signals were sampled at 2 kHz, amplified 500x, and band-pass filtered between 1.0 Hz and 500 Hz. Off-line data analyses were performed by a blinded experienced experimenter to manually detect electrographic seizures utilizing a modified Racine scale. Data from EEG analysis were taken on the 6th day of each of the 1-week recordings, analyzed at 24 h, 2 h in the day and 2 h at night for each week, and after filtering, spikes were detected using custom MATLAB scripts for spike detection.^51,52^ Frequency analyses were performed using fast Fourier transformation (µV^2^/Hz) using LabChart (ADI Instruments) as described previously^51,52^ for total power analysis [delta (0.1–4 Hz), theta (4.01–8 Hz), alpha (8.01–12 Hz), beta (12.01–30 Hz), and gamma (30.01–100 Hz)], and normalized to baseline recordings.

### 2.5 Neurocognitive and behavioral characterization of NKH mouse

Cognitive deficits and hyperactivity are key symptoms of NKH particularly in the attenuated form.^2,6^ Mice were tested for correlative deficits at age 8 weeks within the vulnerable period with no treatment for both B6 and J129 strains and for the J129 strain also under glycine challenge (3% [w/v] in drinking water) beginning 1 week before testing. Detailed testing methods are provided in the supplemental methods. In the Open Field test, mice were placed in a lighted square box and total distance traveled, velocity, and time spent in the center zone versus the periphery recorded with a video tracking software. In the Y Maze Spontaneous Alternation test, used to quantify spatial working memory, testing occurred in a Y-shaped maze with 3 opaque arms at a 120° angle from each other, and the number of arm entries and triads (triad = entry into 3 consecutive arms without revisiting an arm) were recorded to calculate the percentage of correct alternations. Mice normally explore the less recently visited arm, whereas mice with impaired hippocampal function will display fewer correct alternations.^53^ In the Contextual Fear Conditioning test, mice were exposed to 2 mild electric foot shocks in a specific context as described previously.^54-56^ When reintroduced to the same context (without the shock), mice display freezing behavior, indicating contextual-associative learning.^54,57^ This training paradigm was used to assess hippocampal dependent forms of learning in the mice in the form of contextual freezing. In the Radial Arm Water Maze, mice swim in a circular tank with six equally spaced arms radiating from the center, one of which consistently contained an escape platform at the end. Distinct distal visual cues were placed outside the maze for guidance. Over 2 days of testing, the mice learn to rely on the distal visual cues to find the platform. This test examines spatial learning and memory.^58^ Quantitative performance criteria were: time to reach the platform, number of wrong arm entries, and distance swum.

### 2.6 Statistical analysis

Populations are presented as mean and standard deviation (SD) or as median and interquartile range (IQR). Distribution normality was evaluated using Shapiro-Wilk and Kolmogorov-Smirnov statistics. The difference between 2 populations was evaluated by the Student t-test if normally distributed and by a Mann-Whitney-U test if not normally distributed. The difference between multiple populations was evaluated by analysis of variance (ANOVA) with post-hoc Tukey test if normally distributed, or by Friedman’s 2-way ANOVA by ranks with post-hoc pairwise comparisons adjusted by Bonferroni correction for multiple comparisons if not normally distributed. The difference between observed and expected outcomes was evaluated by Chi-square test. Relationships between 2 variables were calculated as the Pearson correlation coefficient for normally distributed populations or the Spearman rank correlation for non-normally distributed populations. Survival was analyzed by Kaplan-Meier statistic with comparisons done by log-rank statistic. For the body weight analysis, for each strain and sex, the weight of the WT mice at each weekly time point followed a normal distribution and at each week the mean and SD were calculated. The body weight of other mouse populations (heterozygote [HET], homozygote mutant [MUT], without treatment or with glycine) were then transformed into a Z-score (weight_weekx_ – mean_weekx of WT mice_)/SD_weekx of WT_. As the Z-scores were normalized for strain, sex, and age, males and females could be combined in a statistical comparative analysis.^59^ Repeated analyses such as EEG recordings, Contextual Fear Conditioning, and Radial Arm Water Maze testing were evaluated using a multifactorial univariate analysis of variance with repeated measures over sequential episodes, and post hoc Tukey HSD test. Statistical significance was set at 0.05. Statistical calculations were done using IBM SPSS Statistics version 29 (IBM, New York, USA), and Statistica™ version 13.5 (StatSoft, Hamburg, Germany).

## 3 RESULTS

### 3.1 Mouse model, genetics, and natural history

The founder mouse was generated from the original E17.1 embryonic stem cells derived from an F1 hybrid of a B6 and a J129 line with the integrating plasmid on the B6 genomic background ensuring integration (Figure 1B). Following injection of the blastocysts, 1 one chimeric mouse was identified, and bred in 2 separate lines for at least 5 generations to either B6 or J129 mice using marker-assisted breeding in a speed congenic breeding strategy (Figure 1E). This allowed the derivation of a strain that was ≥99.8% congenic from the fifth generation. Sanger sequencing of exon 9 verified the presence of the mutation in each strain (Figure 1C), while for breeding purposes, the mutation was verified by a size difference of the polymerase chain reaction product with the mutant allele being larger than the WT due to the remnant lox site (Figure 1D). Using this method, offspring were tested for transmission of the mutation at 2 weeks of age (Table 1). Heterozygote intercrossing of the B6 line showed a significant deviation from the expected distribution for an autosomal recessive condition (Chi-square p=0.0016), an underrepresentation by 36% for the expected MUT group, slightly more in males 43% than in females 31%. In the J129 strain, we also observed a significant deviation from that expected for autosomal recessive inheritance (Chi-square p= 0.05), with a 21% underrepresentation of MUT, 28% in males, and 16% in females. In both strains, there was no deviation from the expected sex ratio (p=0.60 and 0.75, respectively). In the B6 strain within three months, 22/63 male mutant mice died (13 from confirmed hydrocephalus); and 25/62 female mutant mice died (9 from confirmed hydrocephalus). On histology, the hydrocephalus was obstructive at the aqueduct level, similar to the previously reported mutant *Gldc* strains.^24,41,42,60^

**Table 1:** Observed live mice born from heterozygous intercross breedings.

|  | C57Bl/6J |  |  | J129X1/SvJ |  |  |
| --- | --- | --- | --- | --- | --- | --- |
|  | Male | Female | SUM | Male | Female | SUM |
| WT | 84 | 104 | 188 | 157 | 136 | 293 |
| HET | 220 | 181 | 401 | 307 | 299 | 606 |
| MUT | 63 | 62 | 125 | 111 | 125 | 236 |
| p-value | 0.0077 | 0.051 | 0.0016 | 0.0066 | 0.45 | 0.050 |
| Deceased | 22 (35%) | 25 (40%) |  | 0 | 0 |  |
| Hydrocephalus | 13 | 9 |  |  |  |  |

In each strain live mice at 2 weeks of age were genotyped and compared to the expected birth frequencies by Chi-square test to that predicted for an autosomal recessive inheritance pattern with the p-values shown. The number of mice that died in the next three months are reported as deceased. WT = wild-type, HET = heterozygous, MUT = homozygous mutant.

### 3.2 Decreased GLDC enzyme activity and protein levels result in elevated plasma and tissue glycine levels

In planning for the mouse model, we expressed the variants in the mouse *Gldc* gene via transient expression in COS7 cells. This showed that the GLDC p.A394V mutant enzyme had activity of 23.5±2.3 nmol·h^-1^·mg protein^-1^, which was 22.6% of the WT glycine exchange activity of 104±41 nmol·h^-1^·mg protein^-1^, with only a mild reduction in P-protein on Western blot (data not shown). For comparison, in a similar experiment the equivalent human mutation p.A389V showed 11.9% activity of the activity of the WT.^5^

In the mouse, the hepatic residual glycine cleavage enzyme activity in 5-week-old mutant mice was 22±15% in B6 and 16±5% in J129 (p=0.6), whereas in brain, the residual enzyme activity was higher at 30±12% in B6 and 33±2% in J129 (Table S1). Gldc protein levels were decreased as assessed by Western blot. In liver, mutant mice had Gldc protein levels that were 70% of WT for both strains, and in brain regions ranges were: cortex: 42% – 62%, cerebellum: 35% – 33%, hippocampus 24% – 35%, pons: 33% – 50%, and brain stem: 79% – 44% of WT level for B6 and J129, respectively, (all differences p<0.05; **Table S2**). In WT mice, the absolute Gldc protein levels were highest in cortex, cerebellum, and hippocampus and lower in brain stem. Thus, the mutation reduced both enzyme activity and steady state protein levels *in vivo*.

This decreased glycine cleavage enzyme activity resulted in increased glycine levels in plasma and tissues (Table 2). At age 5 weeks, in B6 mice, the plasma glycine levels were doubled in mutant mice (208%) compared to WT, with no difference for heterozygotes, and in J129 mice, plasma glycine levels were also doubled in mutant mice (208%) and increased by 37% in heterozygous mice. Both the WT and mutant plasma glycine levels were a little higher in B6 than in J129 mice, both by ∼30%. In MUT brain tissue, glycine levels increased compared to WT in cortex (186% in J129 and 195% in B6), and in hippocampus (219% in J129 and 190% in B6), with no change in heterozygotes. Glycine levels in all tissues were higher in B6 than in J129 mice. In cerebellum, glycine levels tripled compared to WT (354% in J129 and 324% in B6). In WT mice, plasma glycine levels correlated with hippocampal glycine levels (ρ=0.712, p=0.002) but no other levels correlated. In MUT mice, plasma glycine levels correlated with cerebellar glycine levels r=0.846, CI: 0.572, 0.950, p<0.001), but not with cortex or hippocampal glycine levels.

**Table 2:**
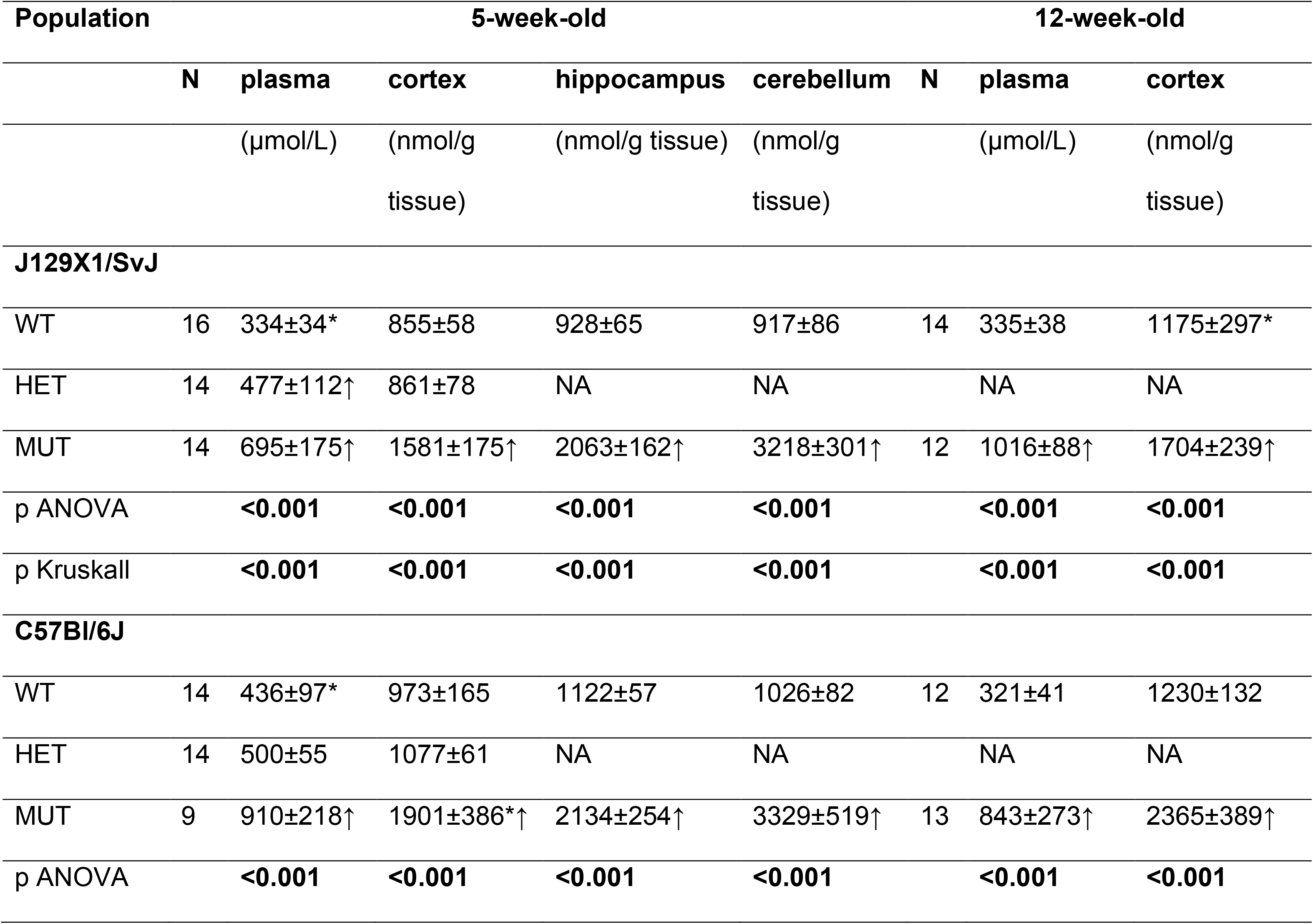
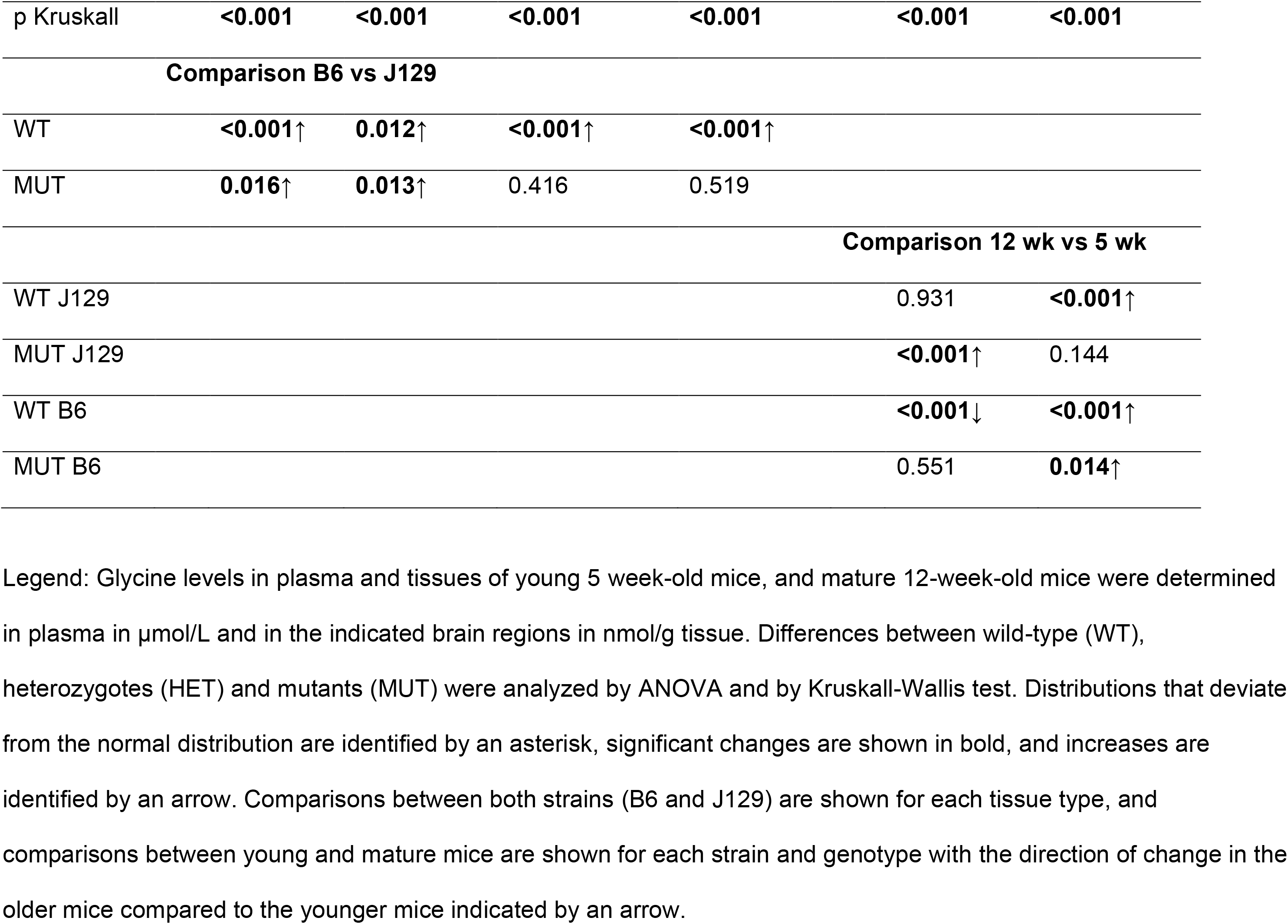
Glycine levels in plasma and brain tissues in young and adult mice.

Compared to young mice at 5 weeks of age, adult mice at 12 weeks had higher plasma glycine levels in J129 (146%) but not in B6, whereas in brain cortex glycine levels were higher in B6 mice (124%) but not in J129. Despite these mild differences by age and strain, mutant mice always had much higher glycine levels in plasma and all brain regions than WT or heterozygous mice. Young mutant J129 mice also exhibited sex differences: glycine levels were higher in female than in male mice in plasma by 45%, cortex by 13%, and cerebellum by 12%; in young B6 mutant female mice glycine levels were higher in plasma by 42% but not in brain levels (Table S3). These sex differences were not present in older mice. In WT mice, there was only a mild sex difference in cerebellar levels in young mice.

When mice were given 5% glycine in drinking water (Table 3), WT mice exhibited markedly increased plasma glycine levels (417% in J129 and 788% in B6), but brain glycine levels increased more moderately in young mice (159% in J129 and 198% in B6) and less or not significantly in older WT mice. In MUT mice, the brain glycine levels increased to 286% in J129 and to 204% in B6. In all cases, MUT mice had about double the cortex glycine levels of WT mice. With glycine challenge in MUT mice, brain glycine levels were 5.3 and 4.0 times the levels in normal WT mouse brain. Thus, increasing the dietary glycine load increased not only plasma glycine levels but also brain glycine levels, more in mutant mice than in WT mice.

**Table 3:** Glycine levels in plasma and brain tissues with glycine challenge.

| Parameter | J129X1/SvJ |  | C57Bl/6J |  |
| --- | --- | --- | --- | --- |
|  | Plasma glycine | Cortex glycine | Plasma glycine | Cortex glycine |
|  | mean±SD (n) | mean±SD (n) | mean±SD (n) | mean±SD (n) |
|  | (μmol/L) | (nmol/g) | (μmol/L) | (nmol/g) |
| **WT** |  |  |  |  |
| 5 weeks water | 334±34* (N=16) | 855±58 (N=16) | 436±97 (N=14) | 973±165 (N=14) |
| 5 weeks glycine | 1395±790*↑ (n=14) | 1356±171↑ (N=14) | 3434±1465↑ (N=14) | 1926±234↑ (N=14) |
| p-value | **<0.001** | **<0.001** | **0.001** | **<0.001** |
| ≥10 weeks water | 335±38 (N=14) | 1175±297* (N=14) | 321±41 (N=13) | 1230±132 (N=13) |
| ≥10 weeks glycine | 1213±662↑ (N=20) | 1214±139 (N=20) | 2322±950↑ (N=10) | 1975±255↑ (N=10) |
| p-value | **<0.001** | 0.604 / 0.051 | **<0.001** | **<0.001** |
| **MUT** |  |  |  |  |
| 5 weeks water | 695±175 (N=14) | 1581±175 (N=14) | 910±218 (N=9) | 1901±386* (N=9) |
| 5 weeks glycine | 8756±5036*↑ (N=14) | 4518±691*↑ (N=14) | 6923±3493↑ (N=16) | 3886±836↑ (N=16) |
| p-value | **<0.001** | **<0.001** | **<0.001** | **<0.001** |
| ≥10 weeks water | 1016±88 (N=12) | 1704±239 (N=12) | 843±273 (N=12) | 2365±389 (N=12) |

|  | New Mouse Model of NKH |  | Swanson et al. |  |
| --- | --- | --- | --- | --- |
| ≥10 weeks glycine | 6916±2464↑ (N=16) | 4037±905↑ (N=16) | 7439±3898↑ (N=12) | 3565±732↑ (N=12) |
| p-value | **<0.001** | **<0.001** | **<0.001** | **<0.001** |
| MUT vs WT on gly |  |  |  |  |
| 5 weeks | **<0.001**↑ | **<0.001**↑ | **0.002**↑ | **<0.001**↑ |
| 10 weeks | **<0.001**↑ | **<0.001**↑ | **<0.001**↑ | **<0.001**↑ |
| on Gly: 10 vs 5 wk |  |  |  |  |
| WT | 0.341 | **0.027**↓ | **0.047**↓ | 0.629 |
| MUT | 0.294 | 0.334 | 0.715 | 0.299 |

Glycine levels in plasma and tissues of young 5 week-old mice, and mature 10 week-old mice were determined in plasma in µmol/L and in brain cortex in nmol/g tissue to evaluate the impact of adding glycine (gly) to the drinking water.
Differences between untreated and glycine treated mice were compared by t-test or Mann-Whitney-U test depending on normality of distribution. Distributions that deviate from the normal distribution are identified by an asterisk, and increases are identified by an arrow, and significant differences shown in bold. Differences between mutant and wild type mice on glycine and between young and older mice on glycine are also provided. WT = wild type, MUT = mutant, Gly = glycine, wk = week.

### 3.3 Natural history and survival under glycine challenge

We next examined the health impact of the mutation on the survival and weight gains of mutant mice. Mutant B6 mice (n=20) had a 40% mortality rate within 90 d, with most deaths occurring between day 35 and 70, primarily due to hydrocephalus, whereas no heterozygous mice (n= 47) and only 1 WT (n=25) mouse was sacrificed for dental malocclusion (p<0.001) (Figure 2A). In the J129 strain, there were no deaths related to NKH in any genotype (WT n= 54, Het n=85, MUT n=23) (Figure 2B). Thus, mortality was higher in B6 than in J129 (p<0.001), but was not different between male and female B6 mice (p=0.88). When B6 mutant mice were challenged with 5% glycine in the drinking water starting at weaning at age 4 weeks, mortality, mostly from hydrocephalus, increased to 60% (n=14), and when challenged from age 1 week, mortality increased to 86% (n=14; p=0.002) (Figure 2C), with no difference by sex (p=0.44).

**Figure 2:**
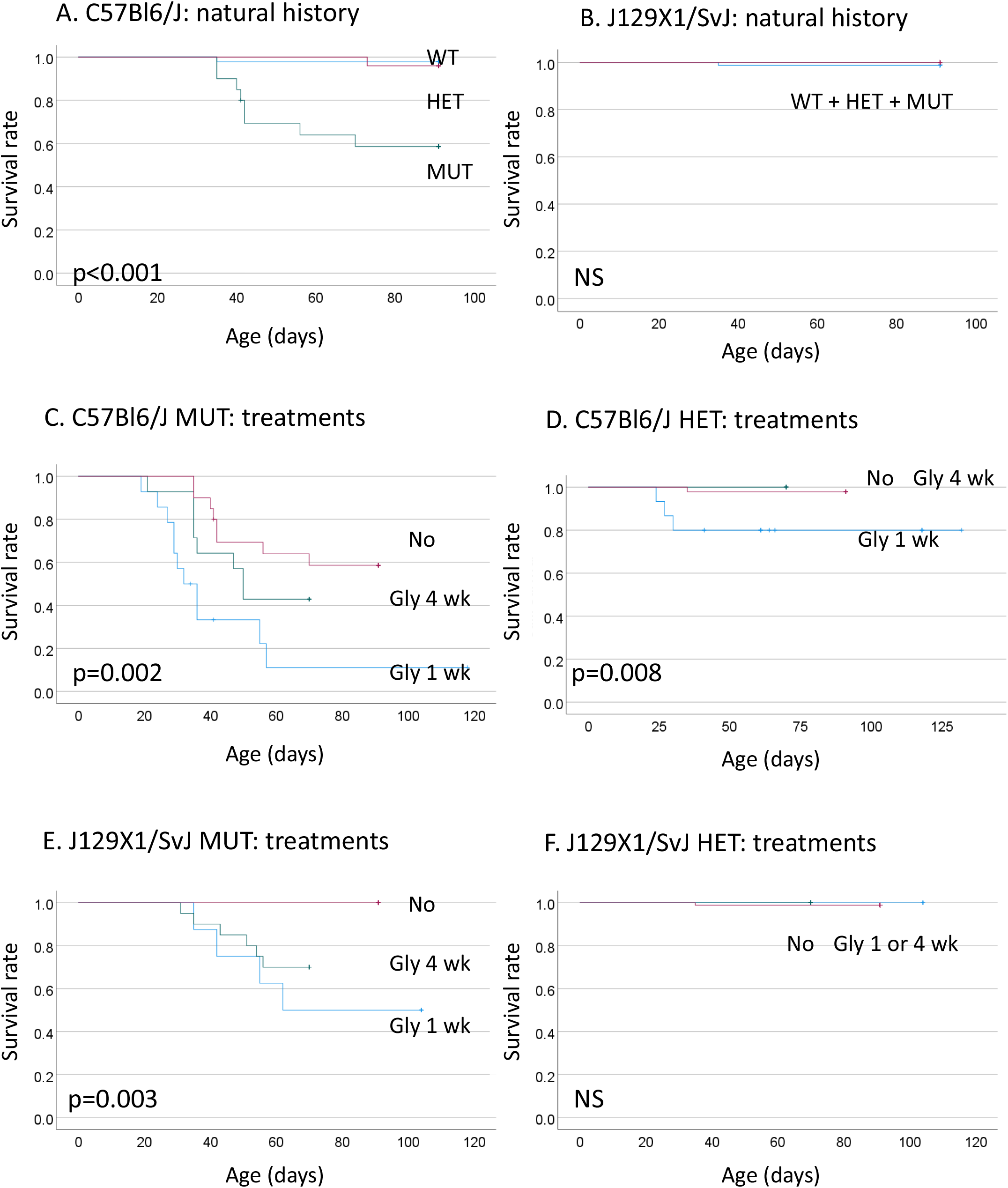
Survival curves for mice with nonketotic hyperglycinemia. **A – B**. The survival curves are shown for mice with wild-type (WT), heterozygotes (HET), and homozygous mutants (MUT) for the C57Bl6/J (**A**) and the J129X1/SvJ (**B**). In B6 mice, MUT (n=20) had a higher mortality than HET (n=47) or WT (n=25), whereas for J129 mice, the MUT (n=23), the HET (n=85) and the WT (n=54) had no mortality. **C – F**. Survival was next compared for mice with no treatment (No) versus challenge with glycine 5% (w/v) in drinking water starting at age 1 week (Gly 1 wk) and after weaning at age 4 weeks on (Gly 4 wk). For B6 mice, the mutant mice had a higher mortality when treated with glycine starting at 1 week (n=14) compared with starting at 4 weeks (n=14) of age and compared with no treatment (n=20) (**C**), whereas heterozygous B6 mice had a mild increase in mortality when treated starting at 1 week (n=15) compared to starting at 4 weeks (n=19) of age and untreated (n=47) (**D**). In J129 mice, mutant mice showed a similar increased mortality when treated starting at 1 week (n=8) or at 4 weeks (n=20) of age compared to untreated (n=23) (**E**). Heterozygotes had no mortality (4 week n=20, controls n=85) (**F**) and neither did WT mice (4 week n=20, untreated n=54). Survival curves for WT treated with glycine are not shown because there was no mortality under any treatment in either strain.

None of the WT mice died, but heterozygotes (n=15) with glycine challenge from 1 week had a 20% mortality compared to none without treatment (n=47) or glycine started at age 4 weeks (n=19; p=0.008) (Figure 2D). In the J129 mice, compared to untreated (n=23), glycine challenge in mutant mice increased mortality to 50%, and was similar when the challenge was initiated at 1 week (n=8) or 4 weeks (n=20; p=0.003) (Figure 2E). In glycine challenged J129 mice, none of the WT or heterozygote mice died (Figure 2F). Thus, glycine challenge increased the mortality of the mutants in both strains. The cause of death in J129 mice was not clear. Mice challenged with glycine at 10 weeks had normal blood count, electrolyte levels, and liver and kidney function. These data indicate a specific postnatal vulnerability for glycine toxicity in the mutant young mice with strain-related differences in severity.

### 3.4 Body weight gain is decreased upon weaning in mutant mice

The body weight of heterozygous and WT mice was similar at all time points in both sexes and strains. In J129 in both sexes, the weight of mutant mice appears to trail that of WT mice in both sexes (Figure 3A,B), albeit less apparent in B6 mice (Figure 3C,D). Mice who develop hydrocephalus had very poor weight gain. On 5% glycine challenge starting after weaning at age 4 weeks weight stagnated in mutant mice compared to WT and HET mice, and was significantly lower for each weekly weight analysis in the J129 (Figure 3E,F) and B6 mice (Figure 3G,H) for both sexes.

**Figure 3:**
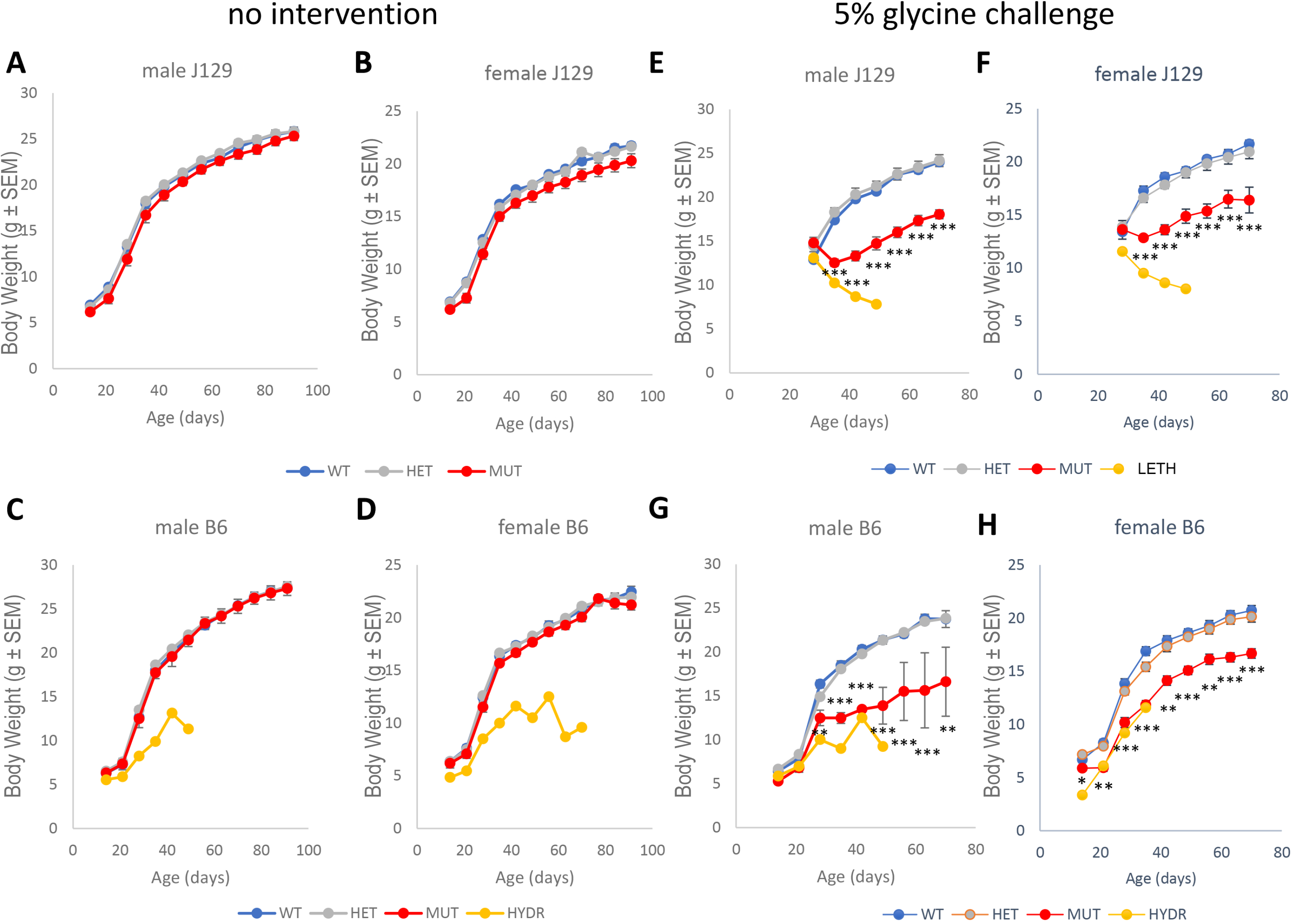
Weight gain in mice with nonketotic hyperglycinemia. The bidy weights of mice are shown as measured every 7 days from age 14 days to 90 days for untreated mice in the natural history (**A-D**) and for mice treated with glycine (5% w/v) in drinking water from days 28 to 72 (**E-H**). The weights of mutant mice without hydrocephalus (MUT, red) are shown in comparison to wild-type (WT, blue) and heterozygous mice (HET, grey), and mutant animals developing hydrocephalus (HYD, yellow) or other early lethality (LETH, in yellow). Mice are shown by natural history without intervention for male (A) and female (B) J129 mice and for male (**C**) and female (**D**) B6 mice. Results are shown following treatment with glycine (5% w/v) in drinking water from week 4 on after weaning for male (E) and female (F) J129 mice and for male (**G**) and female (**H**) B6 mice. Mean±SEM is shown with significance between MUT and WT animals indicated as * p<0.05, ** p<0.01, *** p<0.001.

After transforming the weight at each week for each sex and strain to matched Z-scores, the growth delays become evident. When both sexes were combined, in J129 the weights were reduced in mutant mice compared to WT starting from the earliest time, and continued after weaning but gradually the MUT mice caught up with WT mice in weight, and the difference between WT and MUT was lost by age 8 weeks (Figure 4A). In B6 mice, prior to weaning the mean weight tended to be lower in mutant compared to WT mice, but the difference was not statistically significant. After weaning at 4 weeks, the mutant mice had significantly lower weight compared to WT and HET mice, but regained the weight and caught up to the WT mice by 8 weeks of age (Figure 4B). This catch-up growth was more pronounced in male compared to female mutant mice in both strains (Figure S1). These studies illustrate a vulnerable period of weight delay for 5 weeks following weaning, more notable in J129 than in B6 mice, with a most pronounced effect around 5 weeks of age. When comparing the weight at age 5 weeks, in WT and HET animals, there was no impact of glycine challenge, but in MUT animals in both sexes and in both strains, glycine-treated mice had lower body weight, and further in B6 mice with hydrocephalus the weight was even lower (Figure S2). These data further indicate that the glycine related toxicity in mutant young mice extends to weight gain as well and confirm the age dependency of these effects.

**Figure 4:**
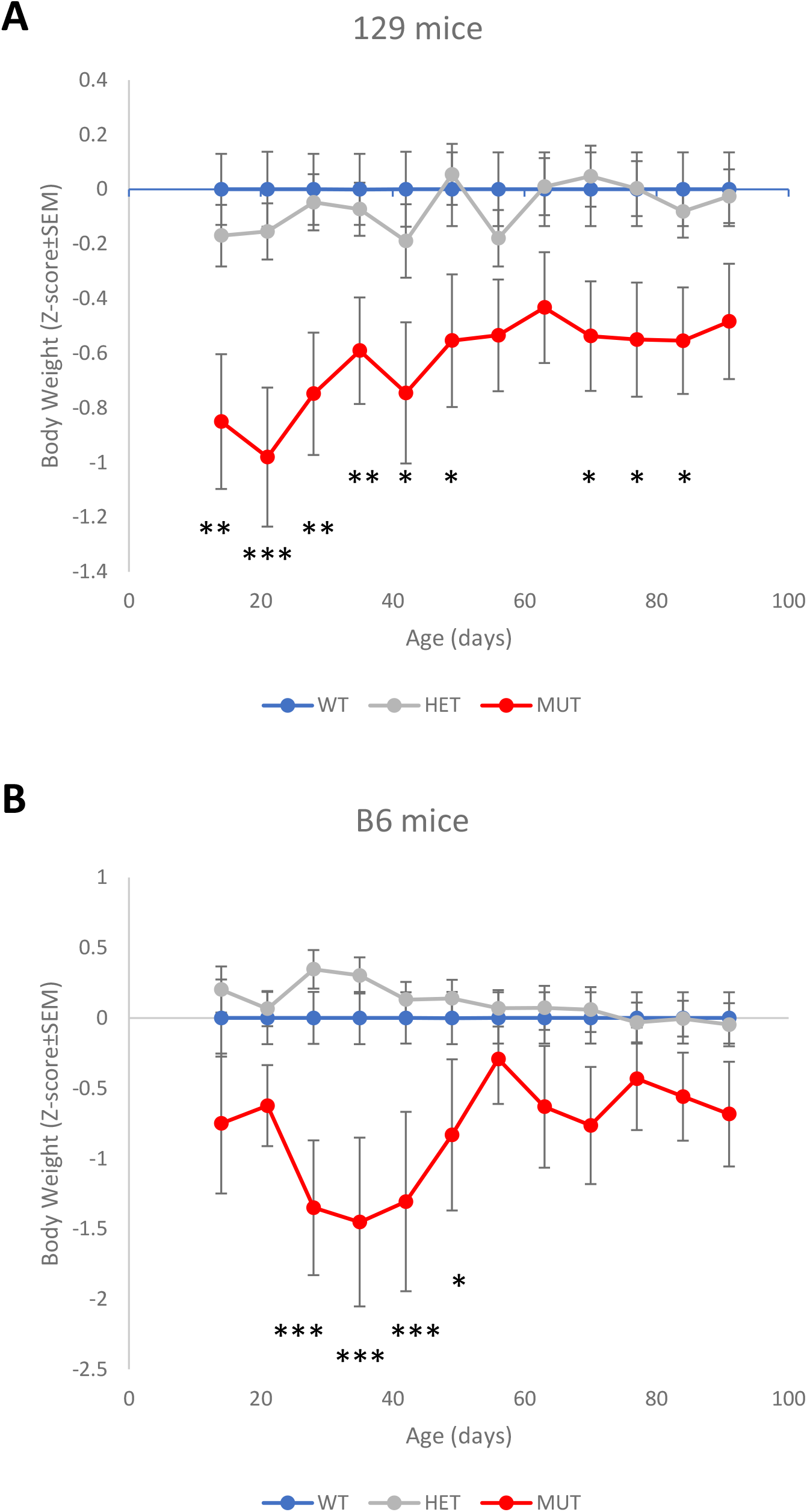
Normalized weight curves in mice with nonketotic hyperglycinemia. The weekly weights of wild type mice followed a normal distribution and the means and SD for each sex and strain calculated. The values of the weights of wild type (WT, blue), heterozygous (HET, grey), and mutant mice (MUT, red) were transformed into Z-scores based on these distributions. For each strain, these sex-adjusted Z-scores were combined for each genotype. **A**. Mutant J129 mice had significantly lower weights compared to controls and heterozygous mice from 3 weeks on and continuing after weaning, but gradually recovered by day 56. **B**. In B6 mice, the youngest mutant mice still had non-significantly lower weights, but after weaning, the weights were significantly lower from ages 28 days to 49 days, but had normalized by age+ 56 days. Mean±SEM is shown with significance between MUT and WT animals indicated as * p<0.05, ** p<0.01, *** p<0.001.

### 3.5 Epilepsy and electroencephalographic recording

Epilepsy and encephalopathy are major symptoms in children affected with NKH, and we evaluated if the MUT mice reflected these symptoms with continuous video-EEG at baseline and glycine challenge. No epileptic events were observed over the 3-week period of continuous video-EEG study in any of the mice. Combining both sexes and strains to compare mutant with WT animals, the spike rate was higher though not significantly in the first week (WT 27.5±4.5 vs. MUT 30.6±8.9, p=0.304) or in the second week (WT 26.3±4.5 vs. MUT 29.8±9.3, p=0.253) without treatment, but in the third week on glycine challenge, the spike rate was significantly higher in mutant animals (WT 25.9±5.1 vs. MUT 35.3±5.1, p=0.011; Table S4). The spike rate intercorrelated between the 3 weeks in the same animals (week 1 vs week 2 Spearman ρ=0.677 p<0.001, and between week 1 and week 3 Spearman ρ=0.652 p<0.001; week 2 versus week 3 Spearman ρ=0.795, p<0.001. This indicates that animals prone to high spike rates do so similarly over the three 1-week recording periods. In MUT mice, the spike rate was significantly higher in week 3 on glycine challenge than in weeks 1 (p=0.002) and 2 (p<0.001). Spike rate was not significantly different by sex or strain.

The spectral band strength for the intensities in the delta, alpha, beta, theta, and gamma frequency ranges were evaluated for a 24-h period, and separately for a 2-h period each during the day and the night. To evaluate for encephalopathic status, we focused on the alpha/delta and the gamma/delta band strength ratio and compared between WT and mutant mice. In the left cortex, the alpha/delta band intensity ratio was significantly lower in mutant than in WT J129 mice during the 24-h period (Table 4), and also separately both in the day and the night for 2-h periods (Table S5). The difference was smaller and not significant in B6 mice. The effect was stronger in female mice, and combining both strains of female mice compared to WT, mutant mice had significantly lower alpha/delta band intensity ratio in the first 2 weeks (WK1 WT 1.02±0.12 vs MUT 0.67±0.12 p<0.001; WK2 WT 1.03±0.18 vs MUT 0.74±0.21 p=0.029) but in the third week on glycine the alpha/delta ratio decreased in both groups resulting in the loss of the difference (WK3 WT 0.75±0.16 vs MUT 0.58±0.19 p=0.123). In multifactorial univariate repeated measures analysis including genotype, sex, strain, and week, the difference by genotype was also significant at p=0.012, with a larger difference in J129 than in B6 (Table 5, Figure 5), with similar findings in the day and the night data (Table S6). While the trend was lower in the right cortex, it did not reach significance (except for week 3 in female mice with both strains combined). In the left hippocampus the alpha/delta band intensity ratio was lower in male and female B6 mice (reaching significance on univariate analysis) and male J129 mice, but surprisingly was higher in mutant female J129 mice, including on univariate analysis (Table 4). This resulted in a significant difference in the genotype by strain combination in the multifactorial analysis for all day (Table 5), and even more strongly for the day and the night recordings separately (Table S6). The right hippocampus mirrored these findings without reaching significance. Indeed, combining all mutant mice a paired sample analysis confirmed the left-right differences in alpha/delta band intensity ratio for the cortex with the left cortex having a significantly lower alpha/delta ratio than the right in the first week WK1 Left 0.81±0.21 vs. Right 0.92±0.28, p=0.05; but lost significance in subsequent weeks (WK2 Left 0.86±0.21 vs Right 0.93±0.32 p=0.38; WK3 Left 0.70±0.21 vs Right 0.72±0.21 p=0.67). The alpha/delta band intensity ratio was not significantly different in the hippocampus. This laterality difference was not present in the WT mice. In the mutant mice, there were significant correlations between both sides in the cortex (WK1 left-right Pearson R^2^=0.793 p=0.002, WK2 0.579 p=0.04, WK3 0.672 p=0.017), but not the hippocampus. The alpha/delta ratio also correlated with the relative strength of the alpha band compared to the combined strength of all bandwidths, but the differences between mutant and WT mice were not significant for the ratio of the alpha band strength out of the total band strengths (Table S7). The gamma/delta band ratio was not significantly different between mutant and WT, and there was no relation between the alpha/delta ratio and the gamma/delta band ratio (data not shown).

**Table 4:**
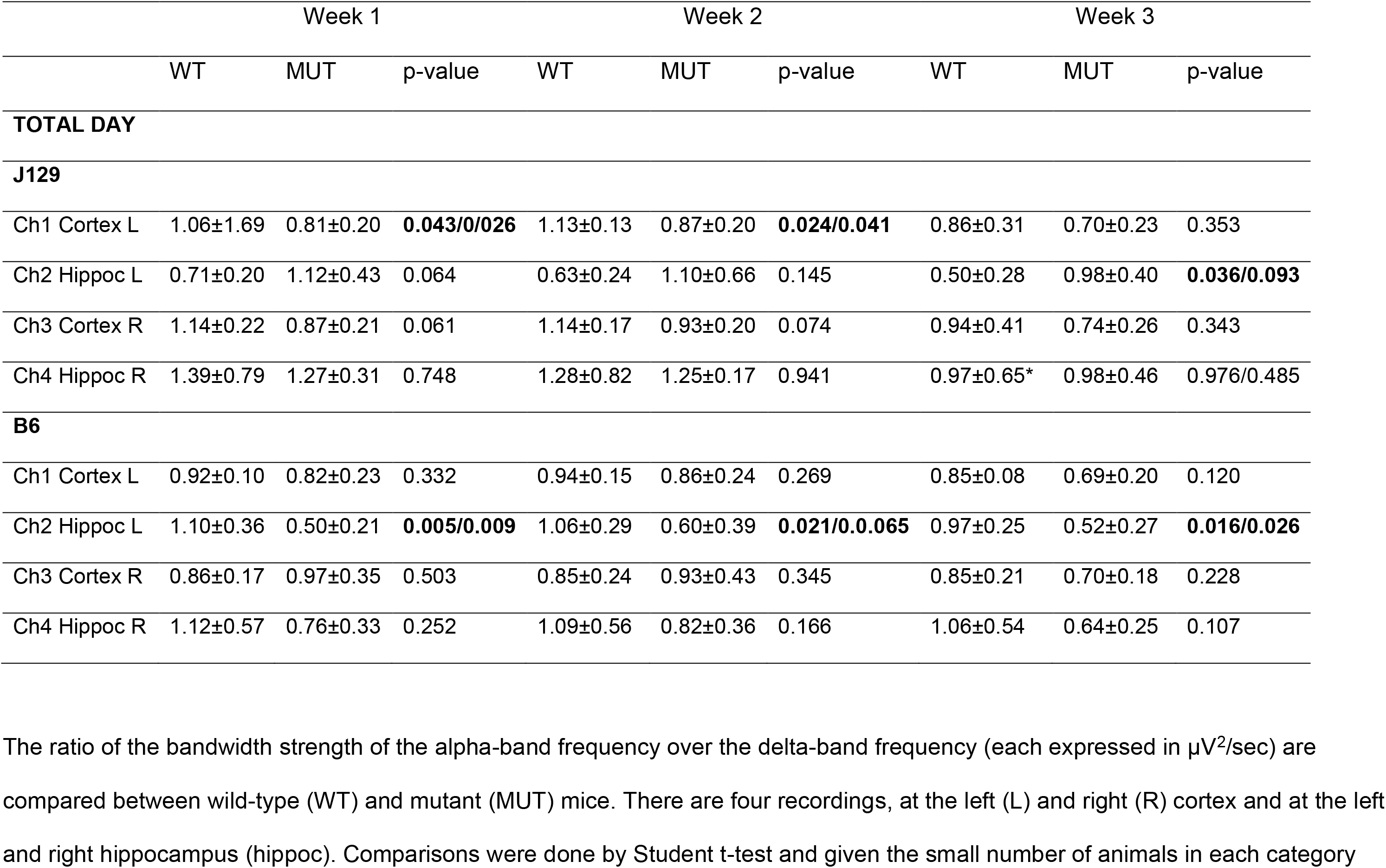
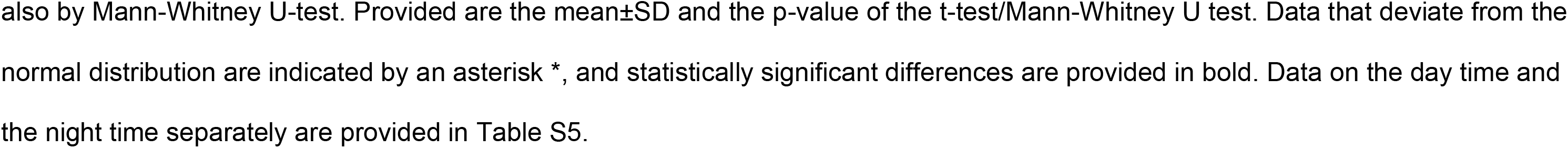
Electroencephalogram power analysis: ratio of alpha/delta spectral band intensities.

**Table 5:** Repeated measures multifactorial univariate analysis of variance of the alpha/delta band signal strength.

| Parameter | Cortex Left |  | Cortex Right |  | Hippocampus Left |  | Hippocampus Right |  |
| --- | --- | --- | --- | --- | --- | --- | --- | --- |
| **ALL DAY** | MS | F, p | MS | F, p | MS | F, p | MS | F, p |
| Genotype | 0.498 | 8.08, **0.012** | 0.198 | 1.23, 0.28 | 0.008 | 0.02, 0.88 | 0.719 | 1.27, 0.28 |
| Sex | 0.399 | 6.48, **0.021** | 0.660 | 4.12, 0.059 | 0.172 | 0.51, 0.49 | 0.672 | 1.19, 0.29 |
| Strain | 0.059 | 0.959, 0.342 | 0.169 | 1.06, 0.32 | 0.046 | 0.14, 0.72 | 1.380 | 2.44, 0.14 |
| Week | 0.188 | 13.61, **<0.001** | 0.193 | 11.04, **<0.001** | 0.101 | 3.18, 0.055 | 0.357 | 10.78, **<0.001** |
| Genotype*sex | 0.185 | 3.01, 0.102 | 0.234 | 1.46, 0.24 | 0.260 | 0.77, 0.39 | 1.800 | 3.18, 0.09 |
| Genotype*Strain | 0.054 | 0.87, 0.36 | 0.260 | 1.63, 0.22 | 4.132 | 12.26, **0.003** | 0.436 | 0.77, 0.39 |
| Sex*Strain | 0.026 | 0.43, 0.52 | 0.001 | 0.00, 0.95 | 0.043 | 0.13, 0.73 | 2.593 | 4.59, **0.048** |
| Geno*Sex*Strain | 0.021 | 0.34, 0.57 | 0.121 | 0.76, 0.40 | 0.301 | 0.89, 0.36 | 0.786 | 1.39, 0.26 |
| Error | 0.06 |  | 0.16 |  | 0.34 |  | 0.56 |  |

Multifactorial analysis of the alpha/delta band signal strength ratio by genotype, sex and strain for the left cortex electrodes analyzed over the entire day. There are significant differences by genotype, and changes by week.

**Figure 5:**
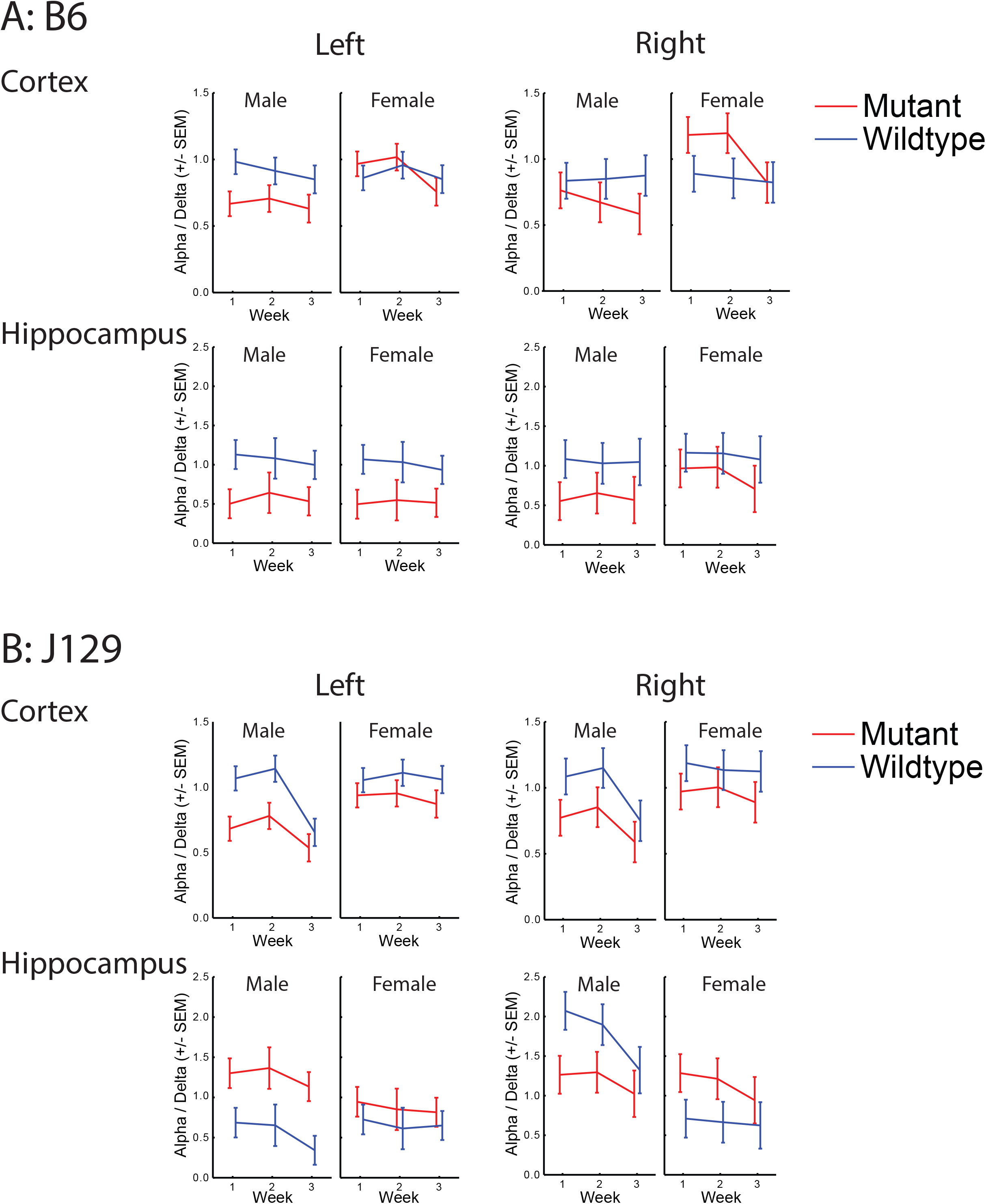
The alpha/delta band intensity ratio for each channel collected over an entire day. The alpha/delta (α/δ) band intensity ratio for each channel was calculated for a 24-h period on day 3 of each week. The mean and standard error for each sex in each strain is given for weeks 1 and 2 when the animal was not treated with glycine, and on week 3 when the animal was supplemented with 5% (w/v) glycine in the drinking water. The EEG was collected over the right and left motor cortex subdural electrodes and in the right and left dorsal CA-1 region of the hippocampus. The WT mice are shown in blue and the mutant mice are shown in red.

In combination, these data indicate a tendency towards increasing spike rate and a propensity for an encephalopathic status with decreased alpha/delta ratio particularly in female J129 mice (which had higher brain glycine levels).

### 3.6 Neurocognitive and behavioral characterization

#### 3.6.1 Open Field Test

The Open Field test evaluates general locomotor activity by recording the distance traveled, velocity and location of the mouse, as well as anxiety-like behavior revealed by lower time spent in the central portion of the box.^61^ On a multifactorial analysis, strain was the dominant factor (F46.9, p<0.001), and B6 WT mice were considerably faster than J129 WT mice (p<0.001; Figure S3A) and traveled almost twice the distance (p<0.001; Figure S3B).

Comparisons in the B6 mice did not identify significant differences for mutation status, but multifactorial comparisons in the J129 mice showed a significant interaction of sex and mutation status (F=4.95, p=0.03; Table S8). Indeed, in J129 males there was no difference in velocity or distance, but the J129 female mutants had significantly lower velocity than WT (6.6±1.5 m/s vs 8.0±1.3 m/s, respectively, p=0.03; Figure S3A), and the distance traveled in MUT was significantly decreased (median 4190 cm [IQR 3784-8603] vs. 7751 cm [IQR 5633-9563], p=0.05; Figure S3B). Under glycine challenge, this difference by genotype in velocity and distance was no longer present. B6 mice spent more time in the center area than J129 (median 241 s (IQR 182-280) vs. 95 s (IQR 45-163) p<0.001; Figure S3C). For B6 mice or untreated J129 mice, there were no statistically significant differences by genotype or sex. For glycine treated J129 mice, male mice spent more time in the center area than females (F=11.181, p=0.003; Figure S3C), and mutants spent more time in the center area than WT, but the difference was not statistically significant (F=4.229, p=0.052).

#### 3.6.2 Y-maze spontaneous alternation test

In the Y-maze, mice explored the 3 arms and the rate of correct alternation was determined based on the first 20 choices (choosing the less recently visited arm). B6 mice were more active than J129 mice as evidenced by a significantly greater number of animals completing the task in the allotted 8 minutes than J129 mice (40/42 B6 mice versus 16/40 J129 mice not treated with glycine (Chi-square p=0.0026) and 17/40 glycine-treated J129 mice). The rate of correct alternations was also higher in B6 than in J129 mice (B6 median 65%, IQR 60-71% vs J129 60%, IQR 51-67, p<0.01). In multifactorial ANOVA, the B6 mice had a small statistically significant difference between WT and MUT in the percentage of correct alternations (F=4.2, p=0.047) with a lower percentage of correct alternations in WT male mice than in the MUT male mice (medians WT = 60%, MUT = 68%, p=0.011; Figure S4A). In J129 mice, there was a strong difference by sex. Female MUT J129 mice explored less with decreased number of arm entries (WT median 20, IQR 16.5-20, MUT median 14, IQR 9.25-20, p=0.013), but no significant difference in the percentage of correct alternations (Figure S4B). Treatment with glycine had no effect on these measures. In males, there was no difference by mutation, but treatment with glycine resulted in a lower percentage of correct alternations (without glycine, median 64%, IQR 60-69.5%, with glycine median 52.9%, IQR 40-65%; Figure S4B,C).

#### 3.6.3 Contextual Fear Conditioning

In this test, freezing can occur in reaction to the foot shocks on day 1, or in anticipation of a shock on day 2. Analysis was performed on 2-min time bins, to allow the characterization of freezing before any shock (Day 1-1), after 2 shocks (Day 1-3), and in anticipation of the shock (Day 2-1). Freezing to the shock was more pronounced in females than males (Figure S5). On a repeated measures ANOVA, there was a significant difference for sex (female more than male, p=0.012) but not of genotype (p=0.224), strain, or glycine challenge. This sex difference was significant in the J129 mice not treated with glycine (Figure S5B, p=0.023). As expected, freezing increased, particularly after the second shock compared to baseline (p<0.01 on Friedman ANOVA). On multifactorial analysis of untreated mice on Day 1 after the first shock, there was a difference by sex (F=7.2, p=0.009) and the difference by mutation status was not significant (F=3.7, p=0.057); on Day1 after the second shock, there were differences by sex (F=11.9, p<0.001) and strain (F=4.9, p=0.03), but not by mutation status. On Day 2, there were differences by sex (p<0.03) in each period with females exhibiting more freezing, but there was no difference by mutation status, or strain. The mutation did not impact the ability of mice to acquire contextual conditioning, pointing to grossly normal hippocampal function.

#### 3.6.4 Radial Arm Water Maze

On the Radial Arm Water Maze, most tested mice were able to learn the position of the hidden platform as shown by the decrease in time to reach the platform in all groups (Figure S6A-C). Six J129 mice that made no effort to reach the platform (floating behavior) were removed from the analyses. The time to reach the platform (Figure S6A-C) and the number of errors were not significantly different by mutation status (Figure S7A-C). For untreated mice, B6 mice performed better than J129 mice, both in time to reach the platform and number of incorrect arm entries (repeated measures ANOVAs, Tukey HSD post-hoc tests p<0.001 both for time to reach the platform and for errors). The difference between WT and MUT mice was not significant overall (repeated measures ANOVA, genotype effect p=0.43) or in either strain (Tukey HSD for time to reach platform: p=0.92 for B6 [Figure S6A], p=0.98 for J129 [Figure S6B); and for number of Errors: p=0.96 for B6 [Figure S7A), p=0.31 for J129 [Figure S7B)).

Glycine challenge in J129 mice had no overall effect on the time to reach the platform (repeated measures ANOVA for Latency, treatment effect p=0.12; for Errors p=0.42; Figures S6C and S7C). Regardless of treatment, female J129 mice performed better than males (Tukey HSD for time to reach platform p=0.009; for errors p=0.44). Specifically for female J129 mice, in the first five trials, the MUT mice did significantly worse for errors (4.90±1.32 vs 3.46±1.30, p=0.012) and a trend for time (41.66±11.64 vs 32.68±9.41, p=0.07). The number of errors and the time to reach the platform intercorrelated for each time point (ρ>0.65, p<0.001). These data further confirm that the mutation does not have an impact on memory function and learning behavior, but does affect the effectiveness in female J129 mice.

## 4 DISCUSSION

To be useful for studying a human condition, mouse models need to recapitulate translational features of the human disease. Such similarities can then be exploited to study pathogenesis or investigate new treatment options. As most treatments in human patients are expected to start after birth, it is important to focus on symptom development after birth that could be impacted by novel treatments. In mice it is imperative to recognize strain and sex differences. Whereas past models only studied NKH models in C57Bl6 mice,^24,25,41,42^ we systematically compared 2 strains and found substantial strain-related differences for symptoms pertinent to NKH such as epilepsy, and motor and cognitive function.^62-64^

A mouse model of NKH with an attenuated variant was generated with reduced enzyme activity *in vivo* and a partial reduction in steady state mutant protein levels, representing a hypomorphic variant. The mechanism for the reduced *in vivo* protein levels remains to be evaluated, with reduced stability of the mutant protein most likely, which could be used to test future potential chaperone molecules. Glycine levels were increased in blood and in brain regions, more pronounced in the cerebellum than in the cortex or hippocampus, similar to what has been observed in postmortem human brain.^14,19,65^ Young female J129 mice had higher brain glycine levels than male J129 mice, which might impact symptomatology. Glycine levels functionally related with development such as growth and brain formation, and with neurophysiologic functioning such as electrophysiology and neurocognitive testing.

Similar to previous studies in B6 mice, fewer mutant mice were born than predicted for an autosomal recessive condition, likely reflecting neural tube defects similar to previous studies.^24,25,42^ The rate of these prenatal defects and the hydrocephalus rate were highly strain dependent, being more prevalent in the B6 than the J129 strain. The B6 strain has a known propensity for hydrocephalus (natural rate of 0.03%),^66,67^ which is exacerbated in all NKH mouse models,^24,25,41,42^ and in other models of folate one-carbon metabolism.^65^ Hydrocephalus also occurs in up to 5% of severely affected human NKH patients,^5,36,68-70^ but neural tube defects have not been reported.^71^ Hydrocephalus was linked to prenatal neural tube defects,^42,60^ and is responsive to prenatal formate treatment.^41,42^ Here we show that hydrocephalus was provoked by postnatal glycine challenge, and that this relates to the timing of the glycine challenge with earlier exposure causing a greater rate of neural tube dysfunction. The symptomatology of neural tube defects occurs in a developmental window between postnatal weeks 4 and 12. The postnatal induction by glycine of these defects indicates that in rodents this process is not a fixed prenatal malformation but rather a fluid developmental condition that can be altered early postnatally, which is a new concept. Whether postnatal treatment may be just as effective as prenatal treatment for the reduction of neural tube defects or hydrocephalus remains to be explored. ^24,25,41,42^

Further, we documented a developmental impact on weight gain in young mice particularly noted after weaning. This developmental impact in young animals reflects the progressive nature of the human disorder, which has the most striking progression of symptoms in the first 2 years of life.^6,72^ Before weaning there appears to be more of an effect in J129 than in B6 mice, which we hypothesize might reflect the poorer rearing capacity of J129 female mice compared to B6.^73^ It is not immediately evident if this growth delay reflects an encephalopathic effect due to increased glycine, or a growth delay related to poor folate methylation. Glycine challenge increased plasma and brain glycine levels more in mutant than in WT mice, resulting in very high brain glycine levels in the mutant mice. The glycine challenge was associated with poor weight gain, in addition to increased mortality without evidence of liver, kidney, or electrolyte dysfunction. The reason for the impact of these levels on the failure to gain weight remains to be explored. In humans not affected by NKH, exposure to high levels of glycine, such as when receiving gamma globulins in a glycine buffer, is not associated with symptoms,^74^ but in NKH patients increasing glycine levels results in lethargy. Surprisingly, glycine-challenged mice did not exhibit decreased locomotor activity in the Open Field test compared to unchallenged mice nor did they develop epilepsy.

Epilepsy is a key symptom of NKH in human patients, which partially improves upon reducing glycine using benzoate therapy or ketogenic diet.^6,28^ While we did not observe signs of epilepsy in our mice, there was an increase in the spontaneous rate of electrographic spikes on EEG, and this increase was significantly exacerbated with glycine challenge. Determination of the epileptic threshold could be a next step. A previous NKH mouse model had more prolonged seizures following an electroconvulsive shock.^38,39^ Cross breeding the mutation to a strain with greater epilepsy propensity (e.g. DBA/2J) may provide a mouse model that better recapitulates this feature in humans.

Human infants with NKH are very encephalopathic and sleepy.^3,6^ Glycine reduction through benzoate or through a ketogenic diet increases wakefulness, thus providing a link to the reduction of glycine to diminish the encephalopathic effect.^26,28,75^ Female J129 mice, who have higher brain cortex glycine levels, appeared to have a lower alpha/delta band intensity ratio on EEG in the left cortex, reflecting a greater encephalopathic effect. Functionally, they moved slower and traveled less in the Open Field test, explored the Y-maze test less, and had worse initial performance on the Radial Arm Water Maze. These similar findings across multiple testing platforms in the same subgroup supports the notion of an encephalopathic phenotype. Thus, a logical next step would be to examine if glycine lowering conditions in these mice improves these functional effects. For a sex-related difference in human NKH patients, an initially reported higher mortality in females^68^ was not replicated in subsequent studies.^5,6,76^

As breeding affected mice resulted in a rapid loss of the more severe phenotype in a previous mouse model of NKH,^37^ we only intercrossed asymptomatic heterozygous carriers. This approach, combined with decreased birth rates and high mortality rates, made it difficult to obtain large cohorts of NKH model mice for the behavioral studies. Therefore, behavioral testing was performed on separate cohorts, sometimes separated by months, which is a limitation of this study.

While our mouse model does not recapitulate all the symptoms of human NKH, it does reflect the biochemistry, the impact on neural tube development such as hydrocephalus, some measures of encephalopathy, and may have a propensity towards seizure-like activity. It does not, however, show microspongiosis and does not develop spasticity and cortical blindness, which are devastating to human patients with severe NKH. Neither do these develop even with a strong glycine challenge, nor have they been reported in any other NKH mouse model. A study of these symptoms may require a different animal model, which is not unprecedented.

For instance, only rat models and not mice of creatine transporter defect and of Lesh-Nyhan disease recapitulate the characteristic neurological symptoms,^77,78^ and in galactosemia the rat, but not the mouse, shows liver dysfunction.^79^ Because the mouse biochemistry reflects that in the human patients, the mouse model can be used to explore therapeutic strategies such as biochemical manipulations, gene therapeutic interventions, or even chaperone therapy given the reduced mutant protein. The in-depth functional characterization of this mouse model will allow investigators to evaluate how such therapies improve functional symptomatology such as measures of brain growth, weight gain, encephalopathic status, and spontaneous spike rate on EEG. The rigorous nature of this animal study controlled for strain effect, age for developmental conditions such as in weight gain, and for sex with an impact on EEG and Open Field test. Such systematic study has allowed us to uncover pertinent symptoms that would otherwise have gone undetected. Moreover, the J129 strain exhibited encephalopathic symptoms not recognized in the B6 strain. They indicate a potential developmental window during which therapies could be most impactful.

Thus, the in-depth phenotyping of this mouse model provides the conditions required to study the impact of new therapeutic interventions for NKH. The variety of symptomatology recognized will allow for the evaluation of which symptoms may relate to specific aspects of biochemical pathophysiology such as elevated glycine or reduced folate one-carbon charging.

## Acknowledgments

We would like to thank the Transgenomics core at the University of Colorado Anschutz campus headed by Dr. Peter Koch for their assistance in generating the transgenic mouse and the NeuroTechnology Center’s Animal Behavior and In Vivo Neurophysiology Core for the behavioral and in vivo neurophysiology studies. We would like to thank Dr. Eric D. Marsh for his input and discussion on the EEG recording and analysis in this manuscript.

## Financial support

Financial support is acknowledged form the NKH Crusaders, Brodyn’s Friends, Nora Jane Almany Foundation, the Dickens Family Foundation, the Lucas John Foundation, Les Petits Bourdons, Joseph’s Fund, the Barnett Family, Maud & Vic Foundation, Lucy’s BEElievers fund, Hope for NKH, Madi’s Mission NKH fund, and from Dr. and Ms. Shaw, and the University of Colorado Foundation NKH research fund. The study was supported by a grant (CNS-X-19-103) from the University of Colorado School of Medicine and the Colorado Clinical Translational Science Institute, which is supported by NIH/NCATS Colorado CTSA Grant Number UM1 TR004399. Contents are the authors’ sole responsibility and do not necessarily represent official NIH views. Funding sources had no role in the design or execution of the study, the interpretation of data, or the writing of the study.

## Conflict of interest

The University of Colorado (JVH, MS, KM, HJ, MWF, RVH) has the intention to file Intellectual property protection for certain biochemical treatments of NKH. Otherwise, the authors have stated that they had no interests which might be perceived as posing a conflict or bias.

## SUPPLEMENTAL INFORMATION

### 1. Supplemental methods

#### 1.1 Mouse Models

Mouse studies were carried out with approval from the Institutional Animal Care and Use Committee of the University of Colorado Anschutz Medical Campus (IACUC# 00413). To generate the transgenic mice, the genomic sequence around exon 9 from the C57BL/6J (B6) mouse strain was cloned into a homologous cloning vector and the p.A394V mutation introduced by site-directed mutagenesis. The vector was transfected for integration using classic techniques into E17.1 embryonic stem cells, which are derived from an F1 hybrid of C57BL/6J and J129X1/SvJ (J129). The phosphoglycerate kinase and neomycin resistance genes flanked by lox sites were removed by transient transfection with Cre recombinase before the cells were injected into blastocysts. Founder mice were backcrossed in 2 separate colonies: B6 and J129 using marker-assisted selection, i.e., a speed congenic approach.

Mouse genomes were assessed at the DartMouse™ Speed Congenic Core Facility at the Geisel School of Medicine at Dartmouth (Lebanon, NH, USA). DartMouse uses the Illumina, Inc (San Diego, CA) Infinium Genotypics Assay to interrogate a custom panel of 5307 single nucleotide polymorphisms (SNPs) spread throughout the genome.^1^ The raw SNP data were analyzed using DartMouse’s SNAP-Map™ and Map-Synth™ software, allowing the determination for each mouse of the genetic background at each SNP location. At each generation a male was selected from 8 to 10 males in a speed-congenics breeding strategy to achieve a congenic status within 5 generations (>99.8% strain background by molecular markers). These 2 separate colonies were used to evaluate the impact of strain backgrounds. The presence of the mutation was verified by Sanger sequencing, and the mutation was traced by polymerase chain reaction (PCR) from tail snip extracted genomic DNA with primers Gldc9-GT-F1: CAGGAGACCTGAACACAGGTGA Gldc9-GT-R1:

ATGCTGGGAACATAACCCACTC, which in the mutant mice generates a longer fragment. The mouse colonies were maintained solely via asymptomatic heterozygote intercrosses to avoid negative selective pressure on the phenotype.

### 1.2 Biochemistry

#### 1.2.1 Enzyme activity measurements

Mutant P-protein activity was measured as percentage of wild-type (WT) in the glycine exchange assay.^2,3^ Initial *in vitro* measurements were performed on COS-7 cells that were transfected with the pCMV6-Entry plasmid containing human WT or p.A389V *GLDC*, or with mouse *Gldc* WT or p.A394V as previously described.^2^For the glycine exchange assay: cell pellets from 150-mm plates were suspended in 600 μL of 20 mM potassium phosphate buffer at pH 7 with 1 mM dithiothreitol (DTT), 1 mM pyridoxal-phosphate, and 1 mM EDTA. Cells were broken by sonication (2 x 20, 1-s pulses at power setting of 5) in a Branson sonicator. Each homogenate was centrifuged at 800x*g* and the supernatant used in the glycine exchange enzyme assay.^3^The assay was carried out by adding 100 µL homogenized sample to 50 mM phosphate buffer Ph 6.2 with 0.1 mM pyridoxal phosphate, 30 mM DTT, lipoylated recombinant human H-protein (approximately 4 µM), 6 mM sodium bicarbonate, and 0.56 mM ^14^C-bicarbonate and the reaction was started by adding 100 mM glycine (150 μL final reaction volume). The mixture was incubated for 30 minutes at 37°C and then glacial acetic acid was added to stop the reaction. After drying and adding scintillation fluid, the radioactivity was measured in a scintillation counter (Beckman LS 6500, Beckman, Indianapolis, IN). Measurements were made in triplicate and mean values from mutant samples were compared to WT controls that were measured on the same day. Activity values were normalized to protein levels determined using the Bradford method.

Enzyme activities were also measured in brain and liver tissues from the mouse model using the glycine cleavage enzyme assay. Approximately 50 mg of tissue was suspended in 500 μL homogenization buffer (20 mM Tris-HCl, pH 8.0, 1 mM DTT, 1 mM pyridoxal phosphate, and 10 µg/ml leupeptin) and homogenized using a ground glass homogenizer. Tissue homogenates (250 μL) or bovine serum albumin blanks (10 μg/mL) were added to: 100 μL homogenization buffer; 25 μL 1 M Tris buffer pH 8; 50 μL THF (25 μg in 6 mL of 100 mM Tris pH 8 with 10 mM beta-mercaptoethanol); 50 μL NAD^+^ (0.66 mg/mL in H_2_O); and 25 μL ^14^C-glycine (0.1 mCi/mL, 55 mCi/mmol) in glass flasks that were fitted with 200 μL potassium hydroxide traps. The reaction was allowed to proceed for 30 minutes at 37°C and was stopped by adding 500 μL of 25% trichloroacetic acid, which also liberated the ^14^C-bicarbonate. The flasks were kept at 37°C for 1 h after adding trichloroacetic acid to allow capture of the ^14^C-bicarbonate in the potassium hydroxide traps. The traps were then added to 5 mL of scintillation fluid (BioSafe II) with 30 μL of glacial acetic acid and were measured in a liquid scintillation counter. Activity values were normalized to tissue mass.

#### 1.2.2 Protein levels

Protein levels were determined using Western blot analysis. In this experiment, 15 J129 mice (7 mutants: 3 males, 4 females and 8 controls: 4 males, 4 females) and 16 B6 mice (8 mutants: 4 males, 4 females and 8 controls: 4 males, 4 females) were used. Liver, cortex, cerebellum, hippocampus, pons, and brainstem were evaluated. Tissues were homogenized as described in the activity assay method above. Tissue supernatant was resolved by sodium dodecyl-sulfate polyacrylamide gel electrophoresis using 10% polyacrylamide gels. Proteins were transferred to Immobilon-P membranes (Millipore), exposed to anti-GLDC antibody (ab97625, Abcam) at 1:5000 dilution, then to goat anti-rabbit secondary antibodies (ThermoScientific, catalog# 31460), and visualized using chemiluminescence (p-coumeric acid, luminol, H_2_O_2_) on X-ray film. Blot images were analyzed using ImageJ (NIH, https://imagej.nih.gov/ij/). Bands from mutants were averaged and compared to bands from WT samples.

#### 1.2.3 Glycine levels

Glycine levels were determined by HPLC in mouse plasma and brain tissue (cortex, cerebellum, and hippocampus) as previously described.^4,5^ In brief, plasma samples were combined with 3:1 volumes of methanol containing 133 µM L-homocysteic acid as an internal standard, and placed on ice for 10 minutes to precipitate the proteins. Brain tissues were homogenized in 2 volumes of phosphate-buffered saline using 0.5 mm zirconium oxide beads in a Storm Pro Bullet Blender (Next Advance, Troy, NY) at speed 4 for 5 minutes, followed by deproteinization as above. Glycine levels were determined after derivatization with o-phthaldialdehyde in combination with N-isobutyryl-L-cysteine using reversed phase HPLC as previously described.

### 1.3 Symptoms, weight, and survival

Mice were maintained on regular chow (Teklad global soy protein-free extruded rodent diet, glycine content 0.7% by weight), with body weight recorded weekly for up to 90 days. Mice that showed signs of severe health distress, usually due to hydrocephalus, were euthanized. Histological sections of a representative brain stained with hematoxylin/eosin were made. To challenge the glycine exposure, glycine (5% w/v) was added to the drinking water supply starting in the first week of life and in another series started at weaning at 4 weeks and continued for 60 days. At the end of each experiment, mice were euthanized by carbon dioxide, plasma was collected, and dissected tissues were flash frozen in liquid nitrogen and stored at −80°C. Samples were handled on dry ice and small portions were removed for subsequent assays. Brain samples were divided into forebrain (cortex), cerebellum, and hippocampus. For subsequent functional studies, mice with hydrocephalus were excluded.

### 1.4 Electrode implantations and EEG recording

Mice were transferred to the electroencephalographic (EEG) procedure room at 4 weeks of age for video-EEG recording.^6-8^ Electrodes were implanted 1 week later. Mice were anesthetized with isoflurane (5% induction, 2-4% maintenance), placed in a stereotaxic frame under normothermic conditions, topical anesthetic was applied subcutaneously, a 1-1.5 cm midline incision was made along the midline of the scalp, the skin was retracted, the skull cleaned, and 0.8-mm burr holes drilled for screw and electrode placements. Subdural electrodes (stainless steel, flat head, self-drilling, 1.19 mm diameter, Fine Science Tools, Foster City, CA, USA) were implanted bilaterally over the sensorimotor cortices [anterior-posterior (AP): −1.0 mm, medial-lateral [ML]: ±3.0 mm relative to bregma) and local field potential depth electrodes (insulated stainless-steel, 0.254 mm diameter, P1 Technologies, Roanoke, VA, USA), were placed in the dorsal hippocampal area CA1 (AP: −2.8 mm, ML: ±2.0 mm from bregma, dorsal-ventral: −2.3 mm from the skull surface), with reference and ground surface electrodes placed over the cerebellum ∼1 mm posterior to lambda, ML ± ∼1.5 mm. The mice were allowed to recover from the surgery for 7 days prior to starting 23 days of continuous video-EEG recordings. For the first 14 days, the mice received regular food and water ad libitum. On day 15, glycine (5% w/v) was added to the drinking water and the mice were recorded for an additional 8 days. Video-EEG monitoring was performed 24 h/day using a digital video-EEG system (Pinnacle Technology Inc, Lawrence, KS, USA) with flexible cables attached to a commutator that allowed the animal to move freely within the recording chamber. EEG signals were sampled at 2 kHz, amplified 500x, band-pass filtered between 1.0 Hz and 500 Hz, and stored on a personal computer.

Off-line data analyses were conducted by a blinded experienced experimenter to manually detect electrographic seizures utilizing a modified Racine scale,^6^ classifying electrographic seizures classes ≥3 as convulsive and classes ≤2 as non-convulsive. Data from EEG analysis were taken on the 6th day of each of the 1-week recordings, analyzed over 24 h, over 2 h during the day and over 2 h during the night) for each week. Prior to data analysis, the data were cleaned by notch-filtering at 60 Hz. Additional EEG analyses were performed using custom MatLab scripts (kindly provided by Dr.

Eric D. Marsh, Children’s Hospital of Philadelphia, Philadelphia, PA) for spike detection as width of 0.1 – 80 ms, and 3 standard deviations above baseline, with those with an absolute amplitude >1200 µV discarded.^9,10^ Frequency analyses were performed using fast Fourier transformation (µV2/Hz) using LabChart (ADI Instruments) with modifications as described^9,10^ for total power analysis [delta (0.1–4 Hz), theta (4.01–8 Hz), alpha (8.01–12 Hz), beta (12.01–30 Hz), and gamma (30.01–100 Hz)]. EEG was normalized to baseline recordings. The spectral band strength for the intensities in the delta, alpha, beta, theta, and gamma frequency ranges was evaluated using a 2-h period during the day from 12 PM to 2 PM and during the night from 12 AM to 2 AM, and over an entire 24-h period. The ratio of the band strength for alpha/delta and for gamma/delta ratios was calculated and compared between WT and mutant mice.

#### 1.4 Neurocognitive and behavioral characterization of the NKH mouse

Cognitive deficits and hyperactivity are key symptoms of NKH particularly in the attenuated form. Mice were tested for correlative deficits at age 8 weeks within the vulnerable period with no treatment in one series for both B6 and J129 strains and in another series for the J129 strain under glycine challenge (3% in drinking water) commenced from 1 week before testing.

#### 1.4.1 Open field

In the open field test, mice were placed in a 44 x 44 cm arena with 25 cm high walls under bright lighting conditions (900-1000 lux) for one 30-min session. Movements and position of the mice were monitored using a video tracking system (Ethovision XT, Noldus, Leesburg, VA). Total distance traveled, velocity, and time spent in the outer zone (≤10 cm from the wall) versus the center zone of the arena were recorded, with the analysis divided in 5-min increments. The open field test is based on the conflicting innate tendencies of mice to avoid bright lights and open spaces thought to mimic a situation of predator risk, and to explore novel environments for increased likelihood of finding food or shelter. The test evaluates both locomotor activity and anxiety-like behavior.^11^ Changes in anxiety were suggested in a previous model of the glycine cleavage enzyme.^12^

#### 1.4.2 Y-maze Alternation test

The Y Maze Spontaneous Alternation test is used to quantify deficits in spatial cognition, whereby normal mice show a tendency to explore the less recently visited arm, resulting in an alternation pattern. Testing occurred in a Y-shaped maze with 3 opaque arms at a 120° angle from each other. After introduction to the center of the maze, the animal was allowed to freely explore the 3 arms. The number of arm entries and the number of triads are recorded in order to calculate the percentage of alternation. A triad was defined as entry into 3 consecutive arms in which the mouse has not re-visited one of the three arms during the triad. An entry occurred when all four limbs are within the arm. Mice had 8 minutes to perform 22 arm entries, which allowed us to calculate 20 possible alternations.

#### 1.4.3 Contextual Fear Conditioning test

This training paradigm used contextual freezing to assess hippocampal dependent forms of learning. Mice were exposed to two mild electric shocks^13-15^ Subsequent exposure without the shock to the same context should elicit freezing behavior, indicating a memory of the adverse experience^13,16^ Mice were placed into 30.5 x 24.1 x 29.2 cm conditioning boxes with a grid floor (rod size: 0.32 cm diameter, rod spacing: 0.79 cm). On the first testing day the mice were placed into the chamber for 6 minutes during which 2 shocks at 2 and 4 minutes were presented after which they were returned to their home cages. On the second day, in the same box, freezing was measured for 6 minutes by a video monitoring system without applying a shock. At least 9 animals were tested for each group (including variables of strain, sex, and genotype (WT or MUT). Freezing behavior during training and testing was assessed using FreezeScan software (CleverSys, Reston, VA), and analyzed in 2-min intervals.

#### 1.4.4 Radial Arm Water Maze

This test utilized a circular swim tank (100-cm diameter) filled with 24±1°C water, with external cues visible to the swimming mice to examine spatial learning and memory.^17^ Six 30-cm long arms equally spaced along the perimeter emanated from the center of the tank. A 12-cm wide platform with a ramp was submerged beneath the surface at the end of one arm. Mice were introduced in another arm in a randomized order. Over 2 days of testing, the mice learn to rely on distal visual cues to find the platform. Mice were introduced in any of the other 5 arms, in a pseudo-randomized order. Training trials comprised 10 swims/d, with a maximum swim duration of 60 s. After 60 s, unsuccessful mice were gently guided to the platform, and stayed there for 10 s. Mice were dried and returned to their home cage, which was kept warm with a heating pad. Quantitative performance criteria were time to reach the platform and number of wrong arm entries. A distinction was made between reference memory errors (entry into an arm that is not the target arm) and working memory errors (re-entry into an arm that has already been visited in the same swimming trial). Some mice can exhibit “floating” behavior, during which they do not try to locate the platform (no or few arm entries within 60 s). Mice were excluded if the number of such trials was above the mean number of occurrences for the entire tested population + 2 standard deviations. Note that the same group of untreated J129 mice were compared to B6 mice and to glycine treated J129 mice. The times from trials 1-5, trials 5-10, and, on the next day, trials 11-15 and 16-20 were combined.

## 3 Supplemental figures

**Figure S1:**
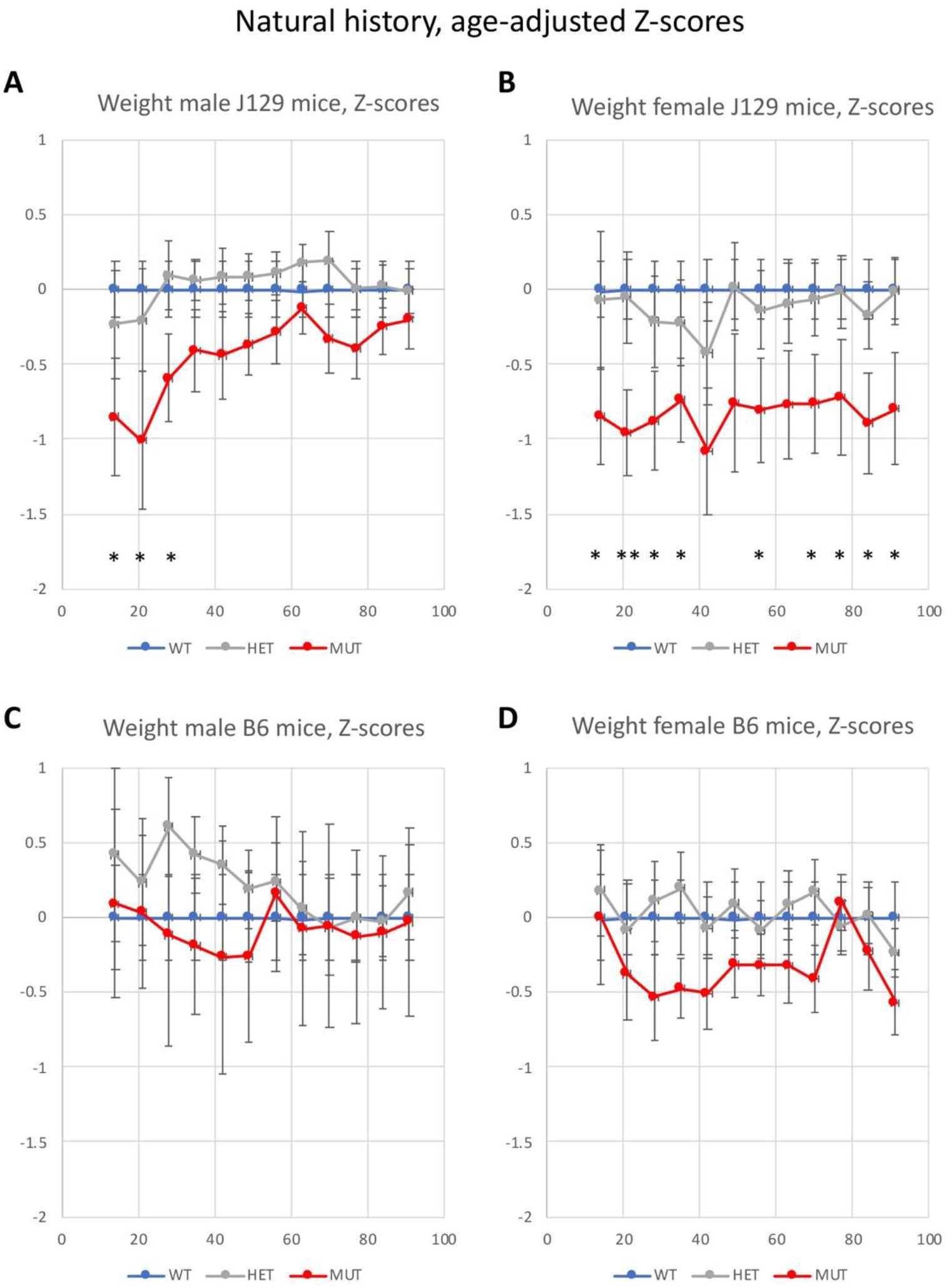
Normalized body weight curves in male and female mice with nonketotic hyperglycinemia. The weekly body weights of wild-type mice followed a normal distribution and the means and SD for each sex and strain were calculated. The values of the weights of wild-type (WT, blue), heterozygous (HET, grey) and mutant mice (MUT, red) were transformed into Z-scores based on these distributions. For each strain, the sex-adjusted Z-scores are shown for each sex for each genotype. The J129 mice had significantly lower weights in male mutant mice compared to controls and heterozygous mice from 3 weeks on and rapidly recovering after 35 days, whereas female J129 mice maintained a lower weight with age. In B6 mice, the youngest mutant mice still had non-significantly lower weights, but after weaning, weights trended lower for 4 weeks in male mice and for a continued period in female mice. Mean±SEM is shown with significance between MUT and WT animals indicated as * p<0.05, ** p<0.01, and *** p<0.001.

**Figure S2:**
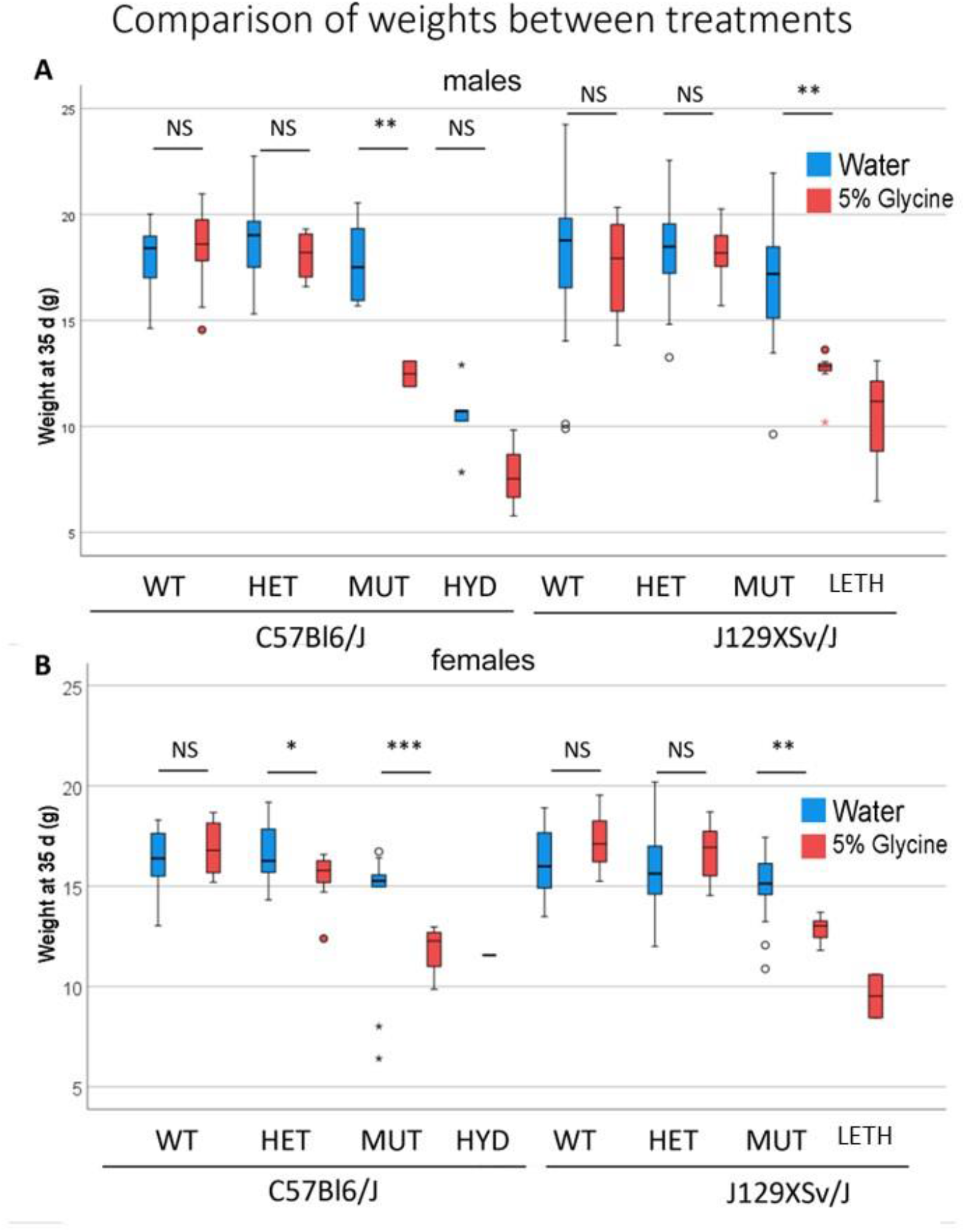
Comparison of the body weight of mice at age 5 weeks by treatment. The body weight of 35-day old mice is shown for C57Bl6/J and for J129Xsv/J mice with male animals in panel A and female animals in panel B. Mice without glycine challenge are shown in blue and mice challenged with 5% glycine in the drinking water are shown in red. Body weight did not significantly differ between treated and non-treated wild-type (WT) and heterozygous (HET) male and female J129 mice, but there was a small decrease in body weight in heterozygous B6 females. Mutant animals (MUT) had a significantly lower body weight at 35 days for both strains and sexes. Mutant animals developing hydrocephalus (HYD) had an even lower body weight. Comparisons were done by Student t-test with * p<0.05, ** p<0.01, and *** p<0.001.

**Figure S3:**
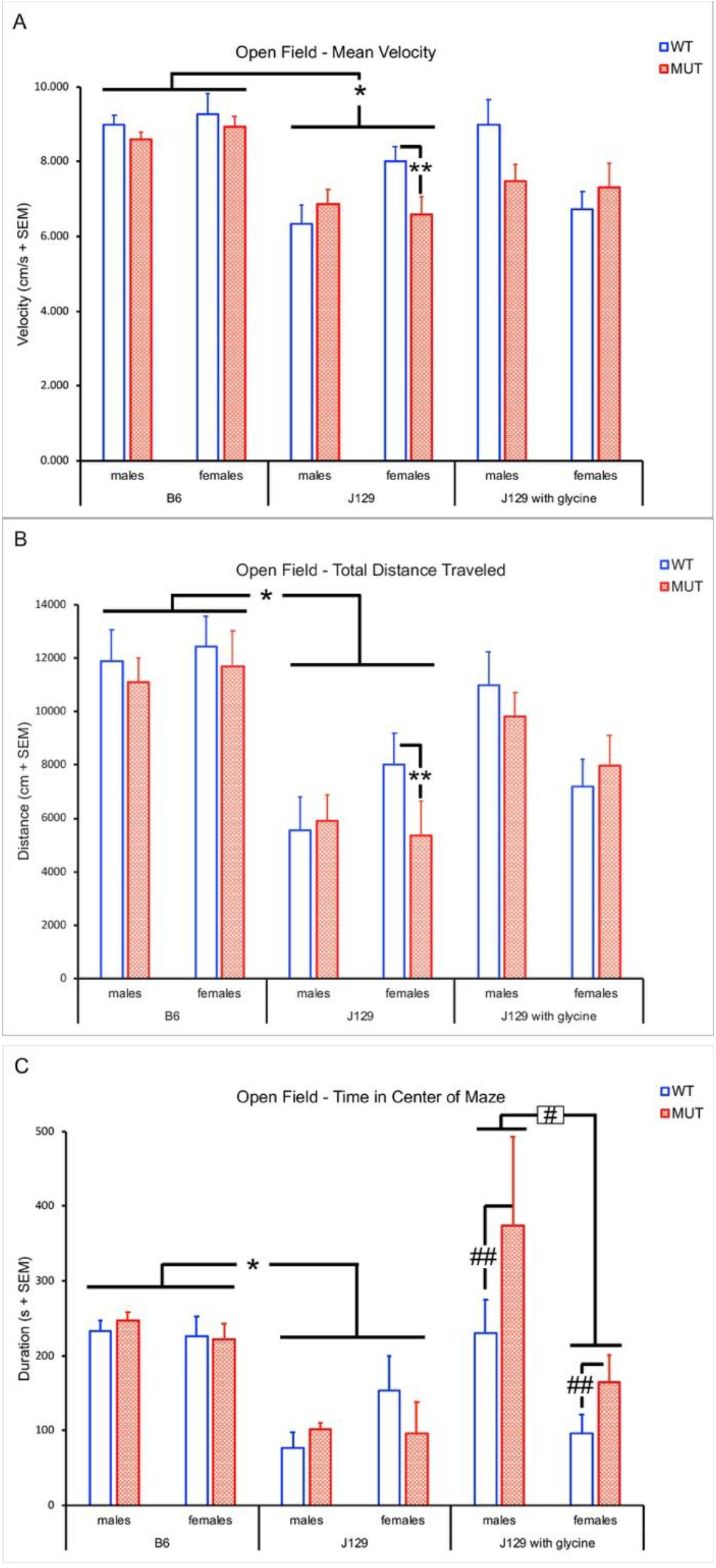
Open Field Test of NKH mice. A. Velocity (cm/s) of WT and MUT mice for both sexes in the B6 and J129 strains. B6 mice had significantly higher velocity than J129 mice. The female MUT J1129 mice had significantly lower velocity than the female WT mice, but there was no significant difference for the male mice between MUT and WT of either strain. B. Distance traveled of WT and MUT mice for both sexes in the B6 and J129 strains. B6 mice travelled significantly further than J129 mice. The female MUT mice travelled significantly less than the female WT mice, and there was no significant difference for the male mice between MUT and WT of either strain. *: p<0.05 between B6 and J129 untreated strains (males and females); ** p<0.05 between female J129 WT and female mutant; # p<0.05 between male and female mice challenged with glycine.

**Figure S4:**
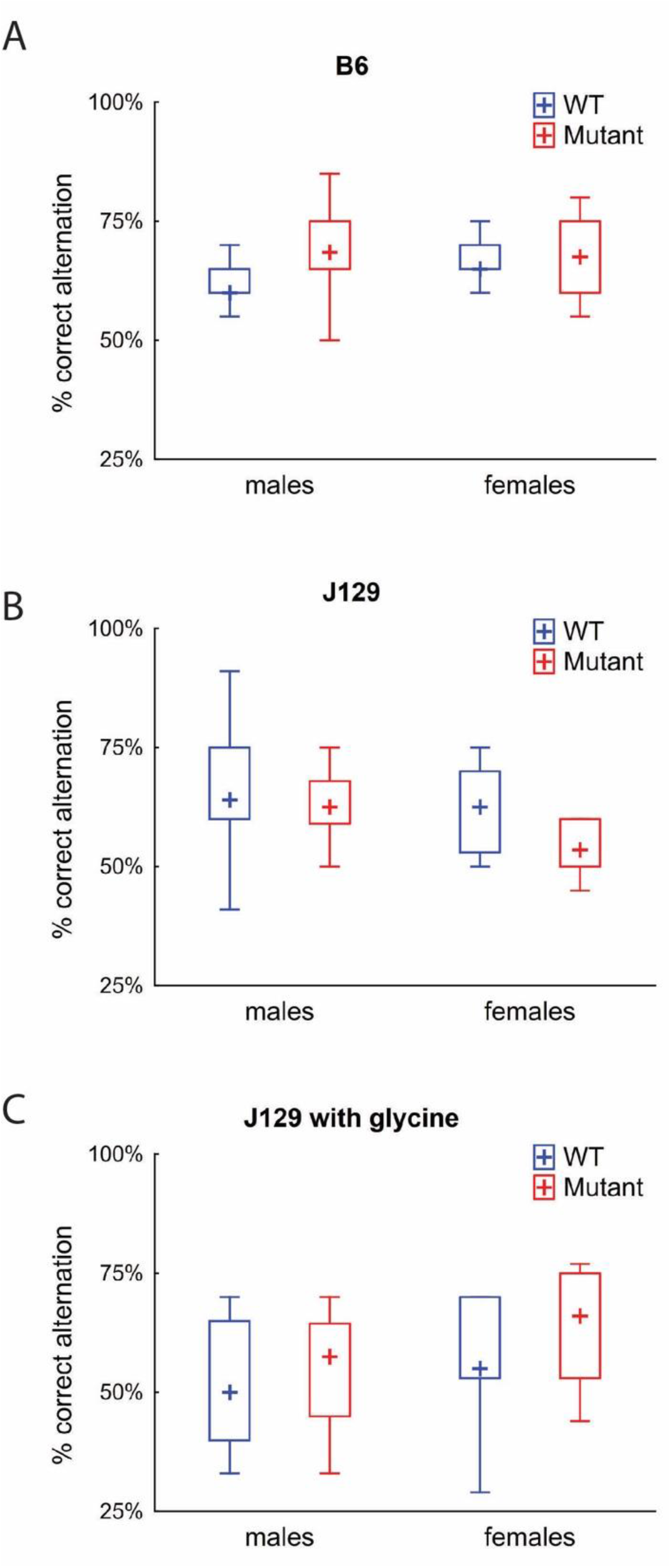
Y-maze Spontaneous Alternation. Percent correct alternations for B6 mice (A), J129 mice (B), and J129 mice challenged with glycine (C). (A). In B6 mice, the WT males make less correct alternations. Multifactorial univariate analysis of variance: model F=2.59, p=0.067; genetics F=4.22 p=0.047, sex F=1.25, p=0.27, genetics*sex F=2.6 p=0.11. (B). For J129 mice there was no significant difference. model F=1.81, p=0.16; genetics F=2.60 p=0.12, sex F=2.51, p=0.12, genetics*sex F=0.32 p=0.58. (C): In J129 mice treated with glycine, there were also no significant differences. model F=1.75, p=0.17; genetics F=1.12 p=0.0.30, sex F=3.86, p=0.057, genetics*sex F=0.04 p=0.84.

**Figure S5:**
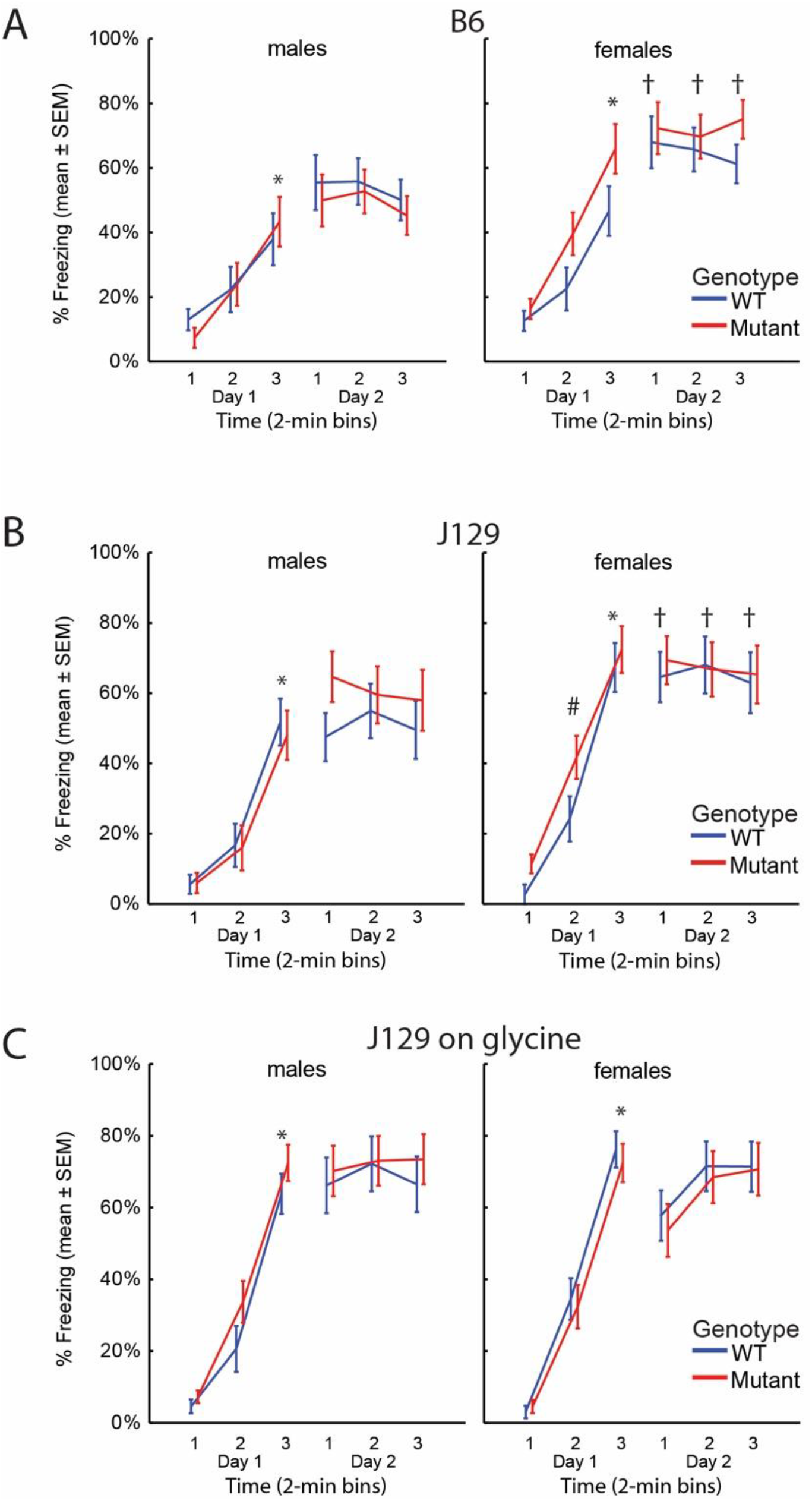
Contextual Fear Conditioning. The contextual fear conditioning test was performed in 39 B6 mice (19 WT [9 male, 10 female] and 20 MUT [10 male and 10 female] mice), 42 J129 mice (21 WT [11 male, 10 female] and 21 MUT [10 male, 11 female]), and 41 glycine challenged J129 mice (20 WT [9 male, 11 female] and 21 MUT [11 male, 10 female]). Freezing was express as the percentage of the time each time, on the first day before treatment (Day1-1), for 2 minutes after the first foot shock (Day1-2), and after the second shock (Day1-3), and for 2 minutes without shock the next day (Day2-1–3). Repeated measures ANOVA showed a difference by time, but not by sex of genotype in each strain. * p<0.05 compared to Day1-1; † p<0.05 from males; # p<0.05 from WT

**Figure S6:**
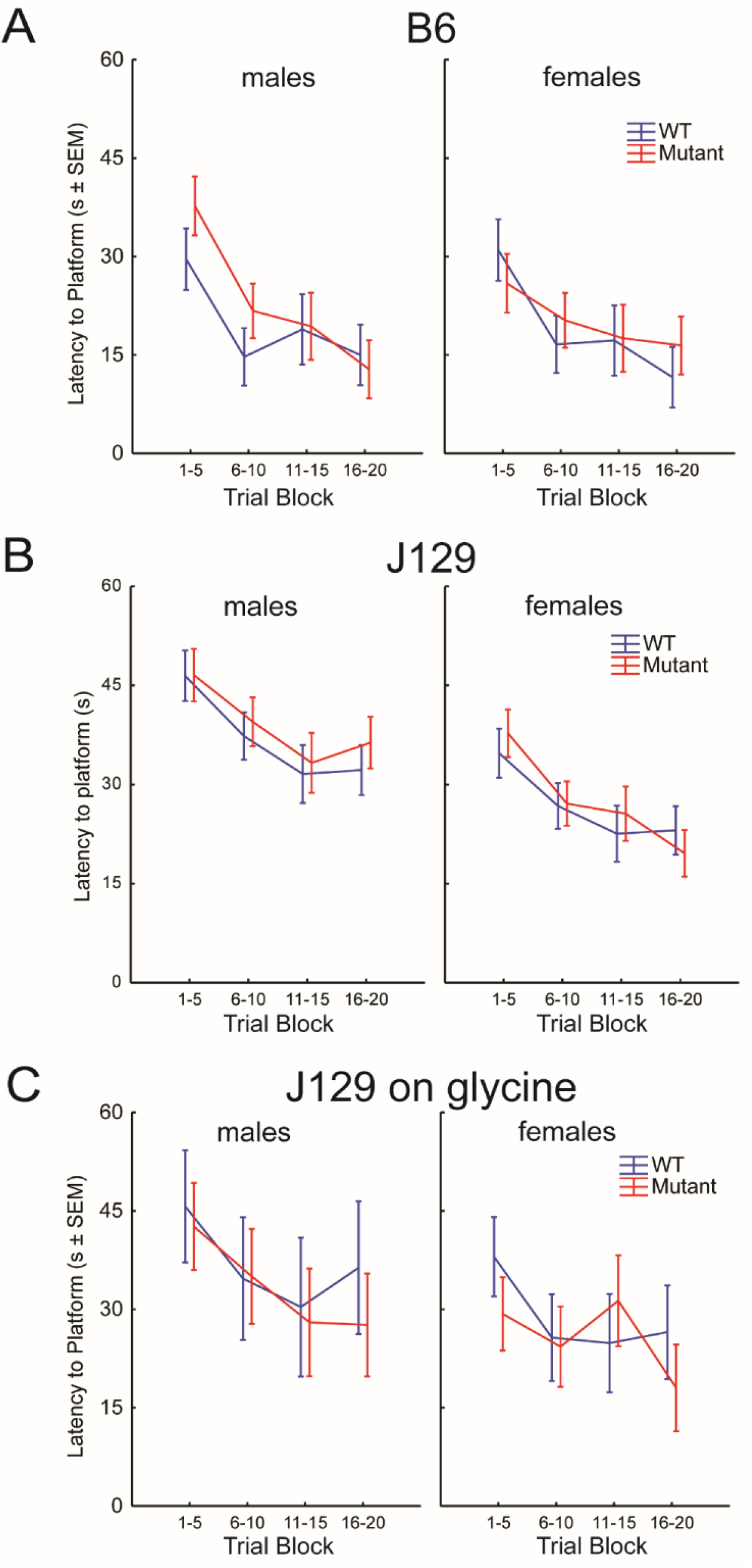
Radial Water Maze: Time to reach the platform. In the Radial Arm Water Maze test, mice are placed in the center of a pool of water and have to find a hidden platform located in 1 of 6 arms. The time to reach the platform (escape latency to platform) for each trial is recorded. Mice perform 10 trials on Day 1, and 10 trials on Day 2, with the mean latency to locate the platform for trials 1-5, 6-10, 11-15, and 16-20 shown. There were 42 B6 mice (10 WT male and 10 WT female and 11 MUT male and 11 MUT female), 44 untreated J129 mice (13 male WT and 10 female WT, 11 MUT male and 10 MUT female), and 24 glycine treated J129 mice (4 WT male and 8 WT female, 5 MUT male and 7 MUT female). A. For B6 mice, on a repeated measures multifactorial analysis, as expected to reflect the learning by the mice the decrease in the time to platform was significant over the time periods F=25.8, p<0.001, but the in between subjects effect was not significant for sex F=0.25, p=0.621, genotype F=0.08, p=0.78, or sex*genotype F=0.05, p=0.82. B. For J129 mice, similarly, the decrease in the time to platform was significant over the time periods F=8.23, p<0.001, and the between subjects effect of sex was significant F=11.38 p=0.002(lower in female), but not by genotype F=1.49, p=0.228 or sex*genotype F=0.19, p=0.66. C. For J129 mice challenged with glycine, decrease in the time to platform was significant over the time periods F=18, p=0.008, but the in between subjects’ effect was not significant for sex F=2.66 p=0.12, genotype F=1.10, p=0.31, or sex*genotype F=0, p=1.0.

**Figure S7:**
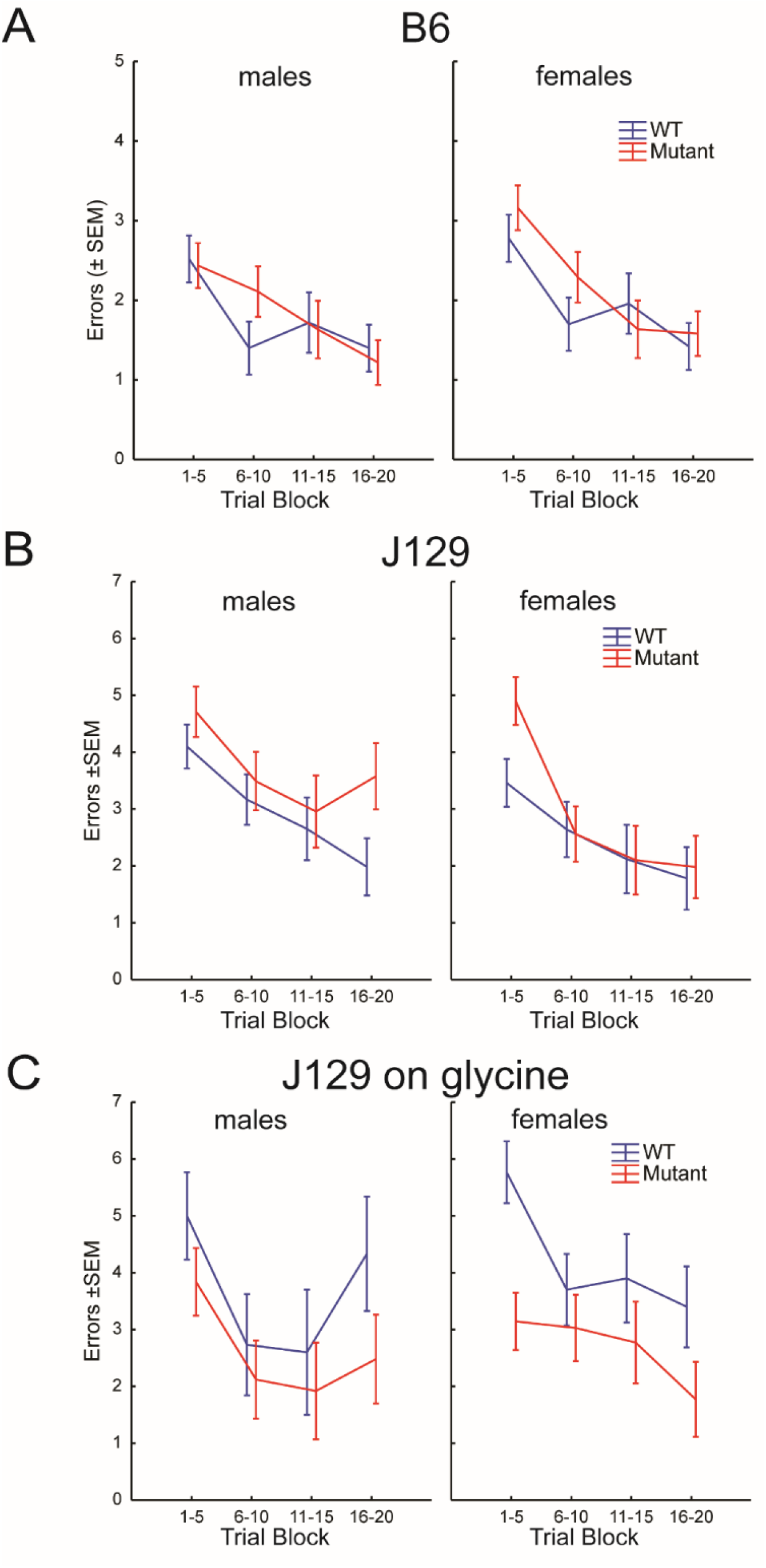
Radial Arm Water Maze: Incorrect Arm Entries. In the Radial Arm Water Maze test, mice are placed in the center of a pool of water and have to find a hidden platform located in 1 of 8 arms. The number of incorrect arm entries (errors) before reaching the platform within 1 minute is recorded. Mice perform 10 trials on Day 1 and 10 trials on Day 2, with the mean number of errors for trials 1-5, 6-10, 11-15, and 16-20 shown. Trials in which the mice make no effort are excluded. There were 42 B6 mice (10 WT male and 10 WT female and 11 MUT male and 11 MUT female), 44 untreated J129 mice (13 male WT and 10 female WT, 11 MUT male and 10 MUT female), and 24 glycine treated J129 mice (4 WT male and 8 WT female, 5 MUT male and 7 MUT female). **A**. For B6 mice, on a repeated measures multifactorial analysis, as expected to reflect the learning by the mice the decrease in the error rate was significant over the time periods F=36, p<0.001, but the in between subjects effect was not significant for sex F=2.16, p=0.15, genotype F=0.67, p=0.42, or sex*genotype F=0.10, p=0.75. **B**. For J129 mice, similarly, the decrease in error rate was significant over the time periods F=38, p<0.001, but not between subjects for sex F=1.89 p=0.18, but not by genotype F=8.86, p=0.27 or sex*genotype F=0.0, p=0.97. **C**. For J129 mice challenged with glycine, decrease in the error rate was significant over the time periods F=18, p<0.001, but and a between subjects effect for genotype F=5.39, p=0.03 (less in MUT), but not for sex F=0.04 p=0.84, or sex*genotype F=0, p=0.97.

## 4 Supplemental Tables

**Table S1:** Residual enzyme activities.

|  | WT | MUT | % |
| --- | --- | --- | --- |
| **LIVER** |  |  |  |
| C57Bl/6J | 6.02 ± 3.08 | 1.32 ± 0.91 | 22% |
| J129X1/SvJ | 4.82 ± 0.35 | 0.76 ± 0.23 | 16% |
| **BRAIN** |  |  |  |
| C57Bl/6J | 1.02 ± 0.37 | 0.31 ± 0.12 | 30% |
| J129X1/SvJ | 1.18 ± 0.38 | 0.38 ± 0.02 | 33% |

Enzyme activity levels of the mouse P-protein were determined using the glycine exchange reaction in triplicate with activities shown in nmol·hour<sup>-1</sup>·mg tissue<sup>-1</sup>.

**Table S2:** P-protein levels.

|  | **C57Bl/6J** |  | **J129X1/SvJ** |  | **WT** |
| --- | --- | --- | --- | --- | --- |
|  | **WT** | **A394V** | **WT** | **A394V** | **[GLDC]** |
| **Tissue** | mean±SD | mean±SD | mean±SD | mean±SD | ng/mg tissue |
| **Liver** | 100±14.8% | 75.8±13.0% | 100±21.1% | 66.6±10.9% | 60-70 |
| **Cortex** | 100±14.5% | 42.6±10.1% | 100±11.7% | 55.3±6.4% | 5-10 |
| **Cerebellum** | 100±13.0% | 34.6±9.6% | 100±9.3% | 32.6±6.4% | 5-10 |
| **Hippocampus** | 100±21.4% | 25.9±12.7% | 100±11.6% | 34.8±5.6% | 5-10 |
| **Pons** | 100±26.9% | 42.6±17.3% | 100±20.1% | 53.3±13.8% | 1-5 |
| **Brain Stem** | 100±17.5% | 76.1±9.8% | 100±23.3% | 41.6±21.1% | 1-5 |

The amount of P-protein quantified on western blot is compared between wild-type (WT) and mutant (MUT) mice for both strains in liver and in different brain regions.

**Table S3:** Glycine levels by sex in untreated mice.

|  | MUTANT |  |  |  | WILD -YPE |  |  |  |
| --- | --- | --- | --- | --- | --- | --- | --- | --- |
| | Male | Female | $\Delta$ | p-value | Male | Female | $\Delta$ | p-value |
| **J129: age 5 weeks** |  |  |  |  |  |  |  |  |
| Plasma | 565±117 | 824±117 | +45% | **0.001** | 325±29 | 344±39 | +6% | 0.408* |
| Cortex | 1486±96 | 1675±191 | +13% | **0.026\*** | 840±61 | 874±52 | +4% | 0.256 |
| Hippocampus | 2031±112 | 2094±205 | +3% | 0.484 | 901±55 | 963±64 | +7% | 0.057 |
| Cerebellum | 3029±247 | 3406±227 | +12% | **0.012** | 961±82 | 861±57 | -10% | **0.016** |
| **B6: age 5 weeks** |  |  |  |  |  |  |  |  |
| Plasma | 768±188 | 1089±61 | +42% | **0.014** | 372±50 | 472±100 | +27% | **<0.001\*** |
| Cortex | 2082±452 | 1676±49 | -19% | 0.115 | 1031±207 | 941±141 | -9% | 0.344 |
| Hippocampus | 2204±195 | 2046±320 | -7% | 0.388 | 1046±76 | 1008±42 | -4% | 0.238 |
| Cerebellum | 3089±565 | 3630±274 | +18% | 0.126 | 1043±69 | 1172±45 | +12% | **0.001** |
| **J129 age 12 weeks** |  |  |  |  |  |  |  |  |
| Plasma | 997±69 | 1035±107 | +4% | 0.485 | 351±39 | 318±30 | -9% | 0.103 |
| Cortex | 1836±114 | 1572±266 | -14% | **0.050** | 1325±364 | 1025±79 | -23% | **0.007\*** |
| **B6 age 12 weeks** |  |  |  |  |  |  |  |  |
| Plasma | 716±255 | 971±245 | +36% | 0.240* | 338±31 | 307±45 | -9% | 0.181 |
| Cortex | 2342±565 | 2388±108 | +2% | 0.850 | 1333±106 | 1142±77 | -14% | **0.002** |

Levels of glycine in plasma in micromolar and in brain tissues in nanomoles per gram tissue are compared between male and female species. Mean and standard deviation, and the p-value for the Student t-test if normally distributed or for the Mann-Whitney-U test if not normally distributed indicated by an asterisk \*, with statistically significant differences highlighted in bold.

**Table S4:**
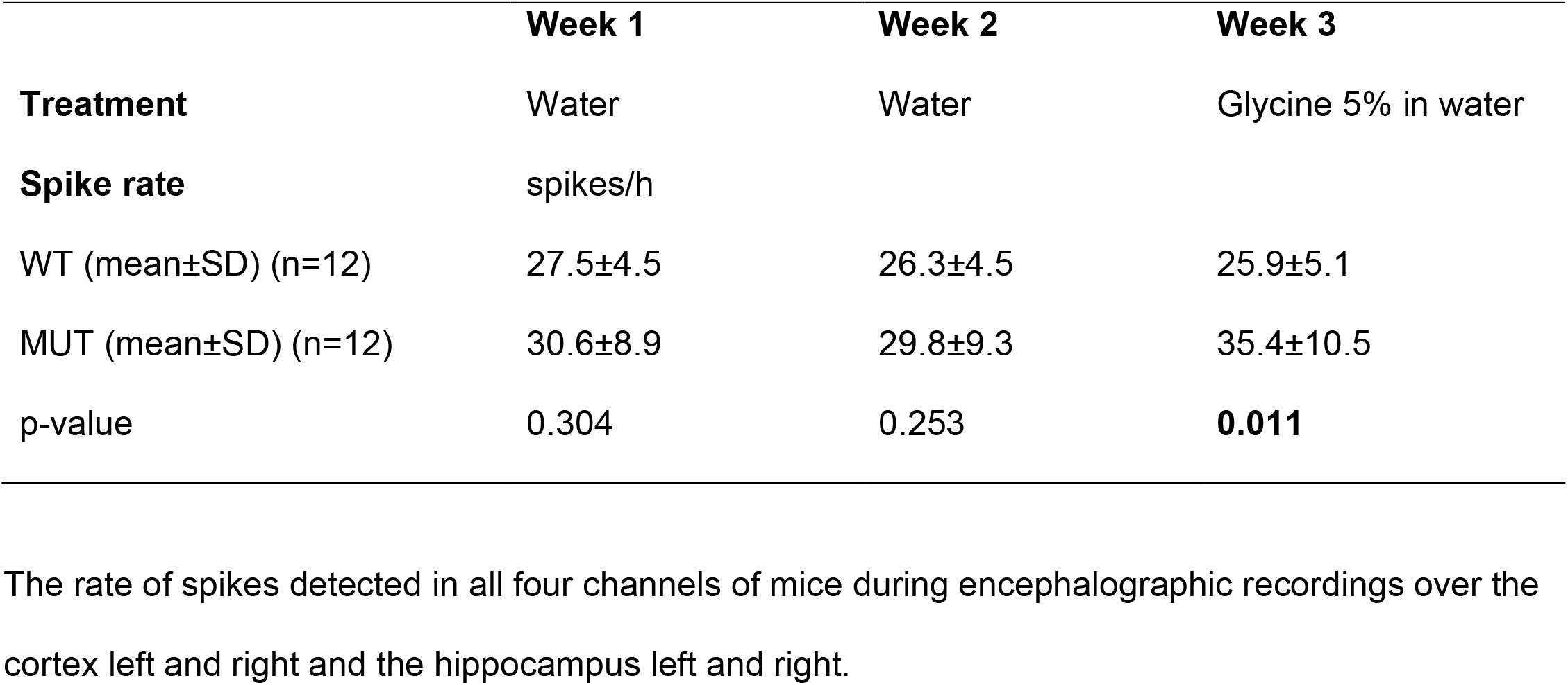
Spike rate in the electroencephalographic recordings of the mice with nonketotic hyperglycinemia.

|  | **Week 1** | **Week 2** | **Week 3** |
| --- | --- | --- | --- |
| **Treatment** | Water | Water | Glycine 5% in water |
| **Spike rate** | spikes/h |  |  |
| WT (mean±SD) (n=12) | 27.5±4.5 | 26.3±4.5 | 25.9±5.1 |
| MUT (mean±SD) (n=12) | 30.6±8.9 | 29.8±9.3 | 35.4±10.5 |
| p-value | 0.304 | 0.253 | **0.011** |

The rate of spikes detected in all four channels of mice during encephalographic recordings over the cortex left and right and the hippocampus left and right.

**Table S5:**
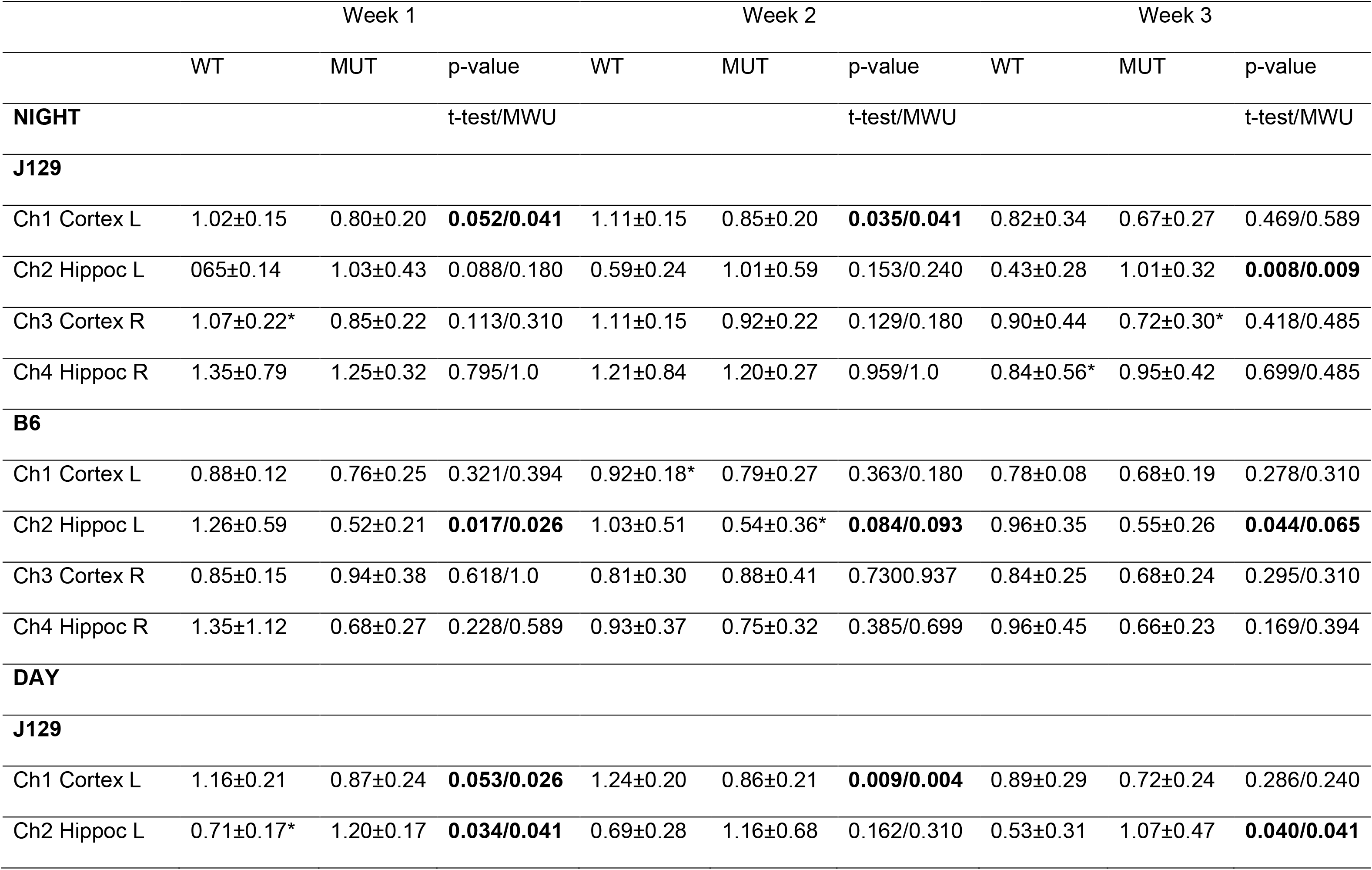
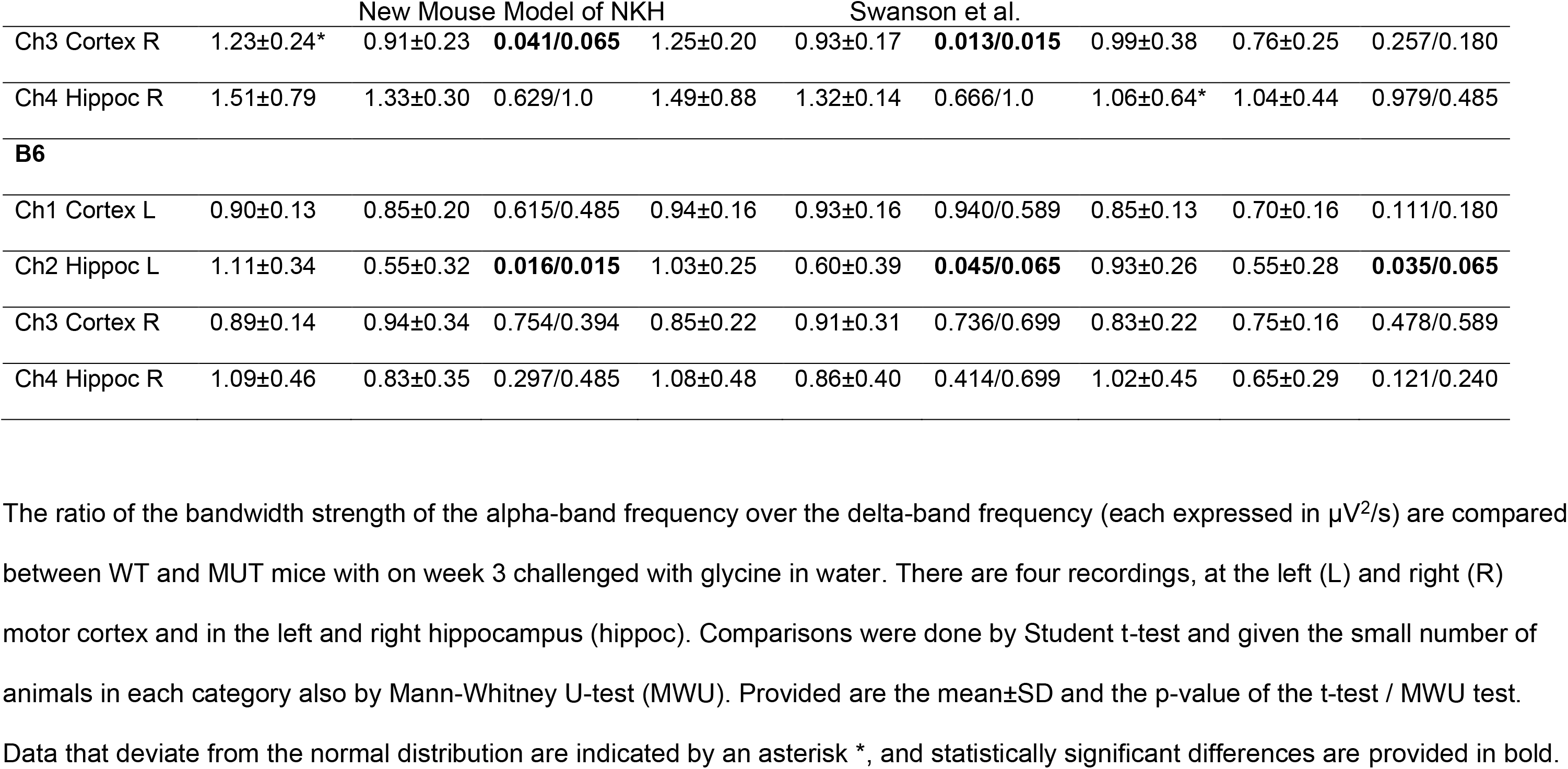

**Table S6:**
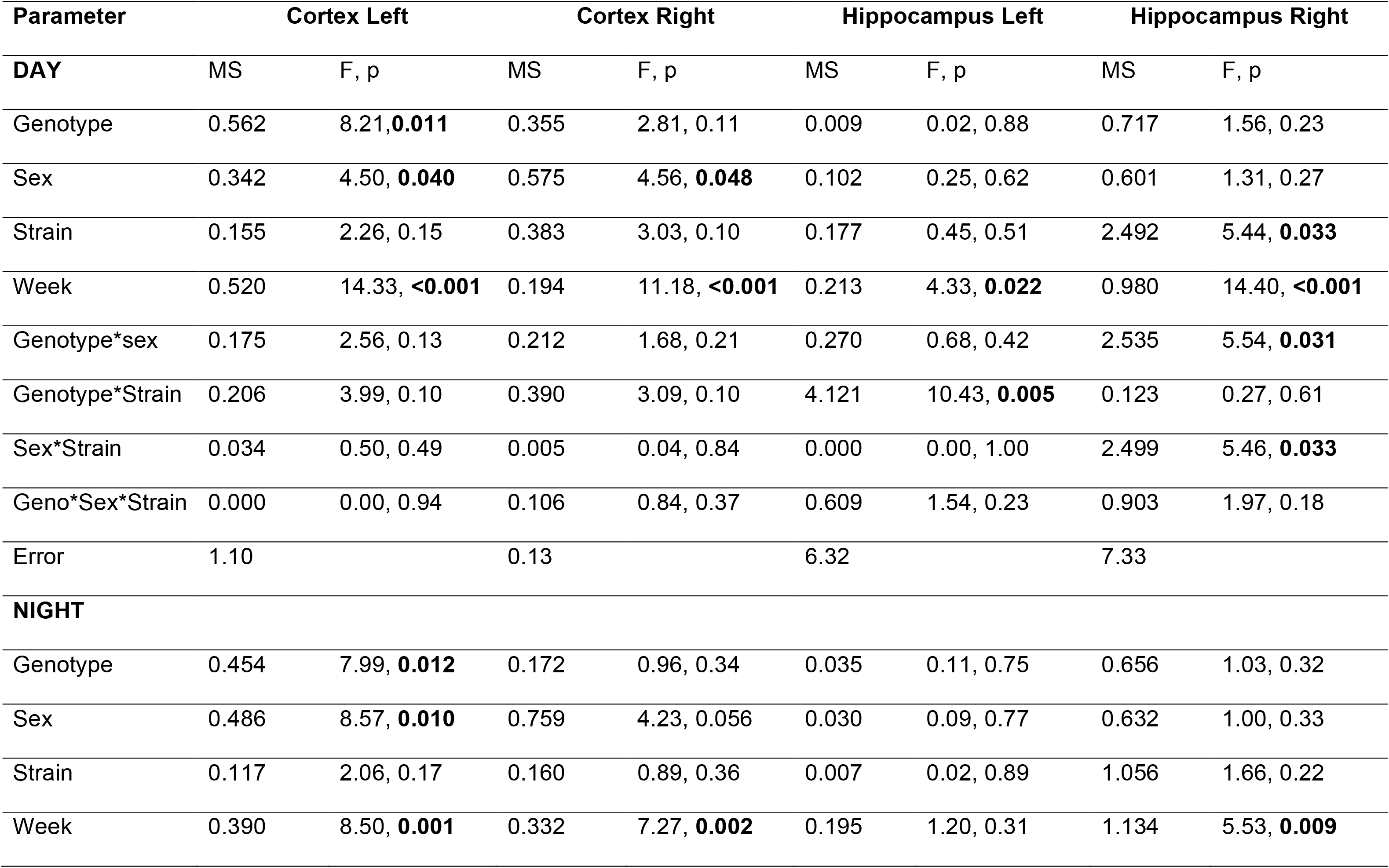
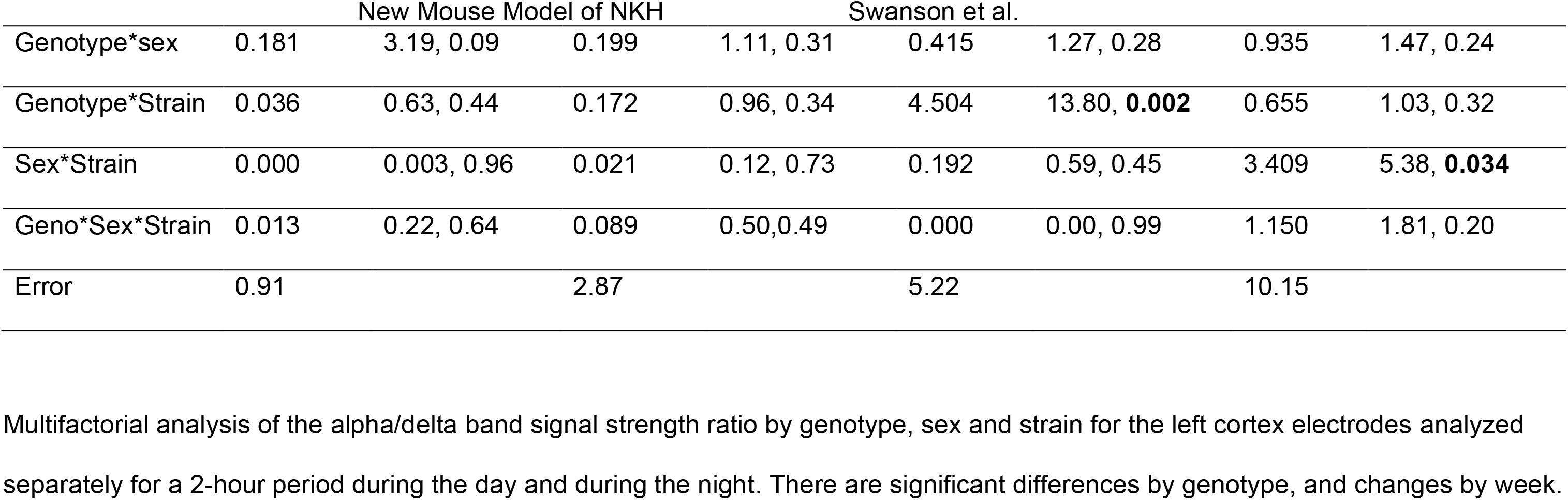
Repeated measures multifactorial univariate analysis of variance of the alpha/delta band signal strength.

**Table S7:**
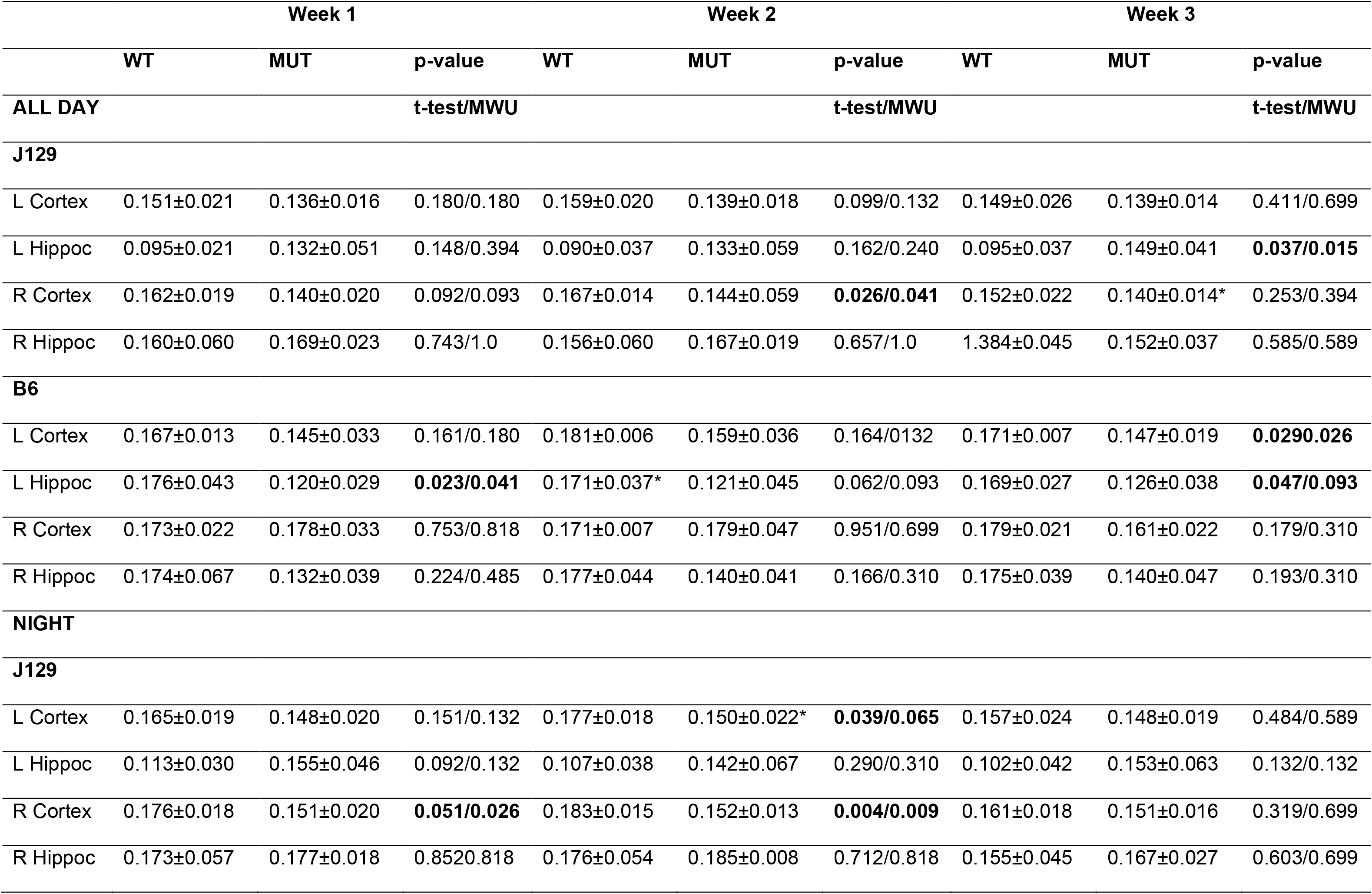
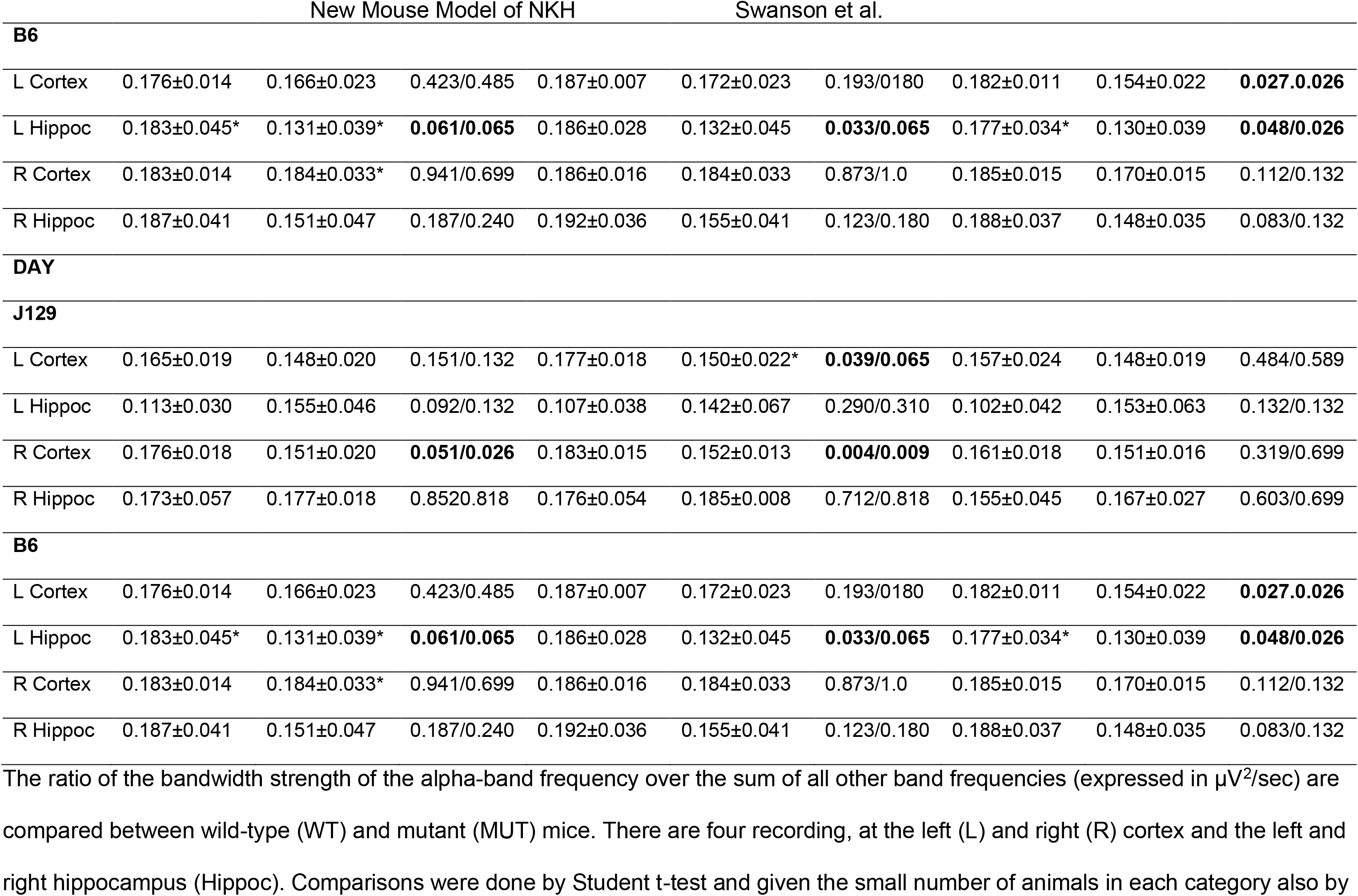
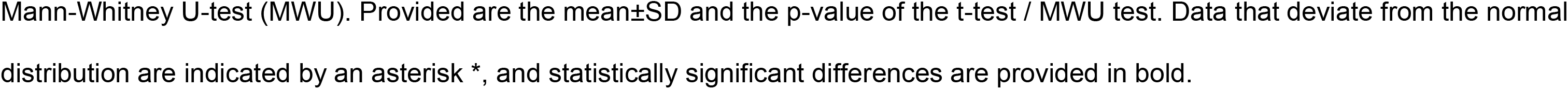
Electroencephalogram power analysis: alpha/total power.

**Table S8:** Open Field Test: Multifactorial analysis of variance for velocity and distance traveled.

A. Velocity
| Dependent variable | df | F | Significance |
| --- | --- | --- | --- |
| Model | 3 | 2.812 | 0.053 |
| Genotype | 1 | 0.979 | 0.329 |
| Sex | 1 | 2.501 | 0.123 |
| Genotype*Sex | 1 | 4.955 | **0.032** |

B. Distance traveled
| Dependent variable | df | F | Significance |
| --- | --- | --- | --- |
| Model | 3 | 2.048 | 0.124 |
| Genotype | 1 | 1.833 | 0.184 |
| Sex | 1 | 1.221 | 0.277 |
| Genotype*Sex | 1 | 3.092 | 0.087 |

Multifactorial analysis using a linear model for the factor velocity showed only statistical significance for the combination of genotype and sex. Note that velocity was calculated only for instances where the animals were moving.

## References

1. Coughlin CR, 2nd, Swanson MA, Kronquist K, et al. The genetic basis of classic nonketotic hyperglycinemia due to mutations in GLDC and AMT. Genet Med. 2017;19(1):104–104. 10.1038/gim.2016.74

2. Van Hove J, Coughlin CI, Swanson M, Hennermann J. Nonketotic Hyperglycinemia. In: Adam M, Ardinger H, Pagon R, et al., eds. GeneReviews® [Internet]. Seattle: University of Washington, Seattle; 2002.

3. Hamosh A, Johnston M. Nonketotic hyperglycinemia. In: Scriver C, Beaudet A, Sly W, et al., eds. The Metabolic and Molecular Bases of Inherited Disease. New York: McGraw-Hill; 2001:2065–2078.

4. Arribas-Carreira L, Dallabona C, Swanson MA, et al. Pathogenic variants in GCSH encoding the moonlighting H-protein cause combined nonketotic hyperglycinemia and lipoate deficiency. Hum Mol Genet. 2023;32(6):917–917. 10.1093/hmg/ddac246

5. Swanson MA, Coughlin CR, Jr., Scharer GH, et al. Biochemical and molecular predictors for prognosis in nonketotic hyperglycinemia. Ann Neurol. 2015;78(4):606–606. 10.1002/ana.24485

6. Hennermann JB, Berger JM, Grieben U, Scharer G, Van Hove JL. Prediction of long-term outcome in glycine encephalopathy: a clinical survey. J Inherit Metab Dis. 2012;35(2):253–253. 10.1007/s10545-011-9398-1

7. Anderson JM. Spongy degeneration in the white matter of the central nervous system in the newborn: pathological findings in three infants, one with hyperglycinaemia. J Neurol Neurosurg Psychiatry. 1969;32(4):328–328. 10.1136/jnnp.32.4.328

8. Shuman RM, Leech RW, Scott CR. The neuropathology of the nonketotic and ketotic hyperglycinemias: three cases. Neurology. 1978;28(2):139–139. 10.1212/wnl.28.2.139

9. Brun A, Börjeson M, Hultberg B, Sjöblad S, Akesson H, E L. Neonatal, non-ketotic hyperglycinemia. A clinical, biochemical and neuropathological study including electron microscopic findings. Neuropädiatrie. 1979;10:195–205.

10. Dalla Bernardino B, Aicardi J, Goutières F, Plouin P. Glycine encephalopathy. Neuropädiatrie. 1979;10:209–225.

11. Agamanolis DP, Potter JL, Herrick MK, Sternberger NH. The neuropathology of glycine encephalopathy: a report of five cases with immunohistochemical and ultrastructural observations. Neurology. 1982;32(9):975–975. 10.1212/wnl.32.9.975

12. Scher MS, Bergman I, Ahdab-Barmada M, Fria T. Neurophysiological and anatomical correlations in neonatal nonketotic hyperglycinemia. Neuropediatrics. 1986;17(3):137–143. 10.1055/s-2008-1052515

13. Rogers T, al-Rayess M, O’Shea P, Ambler MW. Dysplasia of the corpus callosum in identical twins with nonketotic hyperglycinemia. Pediatr Pathol. 1991;11(6):897–897. 10.3109/15513819109065486

14. Agamanolis DP, Potter JL, Lundgren DW. Neonatal glycine encephalopathy: biochemical and neuropathologic findings. Pediatr Neurol. 1993;9(2):140–140. 10.1016/0887-8994(93)90051-d

15. Press GA, Barshop BA, Haas RH, Nyhan WL, Glass RF, Hesselink JR. Abnormalities of the brain in nonketotic hyperglycinemia: MR manifestations. AJNR Am J Neuroradiol. 1989;10(2):315–315.

16. Mohammad SA, Abdelkhalek HS. Nonketotic hyperglycinemia: spectrum of imaging findings with emphasis on diffusion-weighted imaging. Neuroradiology. 2017;59(11):1155–1163. 10.1007/s00234-017-1913-0

17. van der Knaap M, Valk J. Magnetic resonance of myelination and myelin disorders. 3rd ed. Berlin: Springer; 2005.

18. Bekiesiniska-Figatowska M, Rokicki D, Walecki J. MRI in nonketotic hyperglycinaemia: case report. Neuroradiology. 2001;43(9):792–792. 10.1007/s002340100577

19. Perry TL, Urquhart N, MacLean J, et al. Nonketotic hyperglycinemia. Glycine accumulation due to absence of glycerine cleavage in brain. N Engl J Med. 1975;292(24):1269–1269. 10.1056/NEJM197506122922404

20. McDonald JW, Johnston MV. Nonketotic hyperglycinemia: pathophysiological role of NMDA-type excitatory amino acid receptors. Ann Neurol. 1990;27(4):449–449. 10.1002/ana.410270419

21. Tada K, Kure S. Non-ketotic hyperglycinaemia: molecular lesion, diagnosis and pathophysiology. J Inherit Metab Dis. 1993;16(4):691–691. 10.1007/BF00711901

22. Deutsch SI, Rosse RB, Mastropaolo J. Current status of NMDA antagonist interventions in the treatment of nonketotic hyperglycinemia. Clin Neuropharmacol. 1998;21(2):71–71.

23. De Groot CJ, Troelstra JA, Hommes FA. Nonketotic hyperglycinemia: an in vitro study of the glycine-serine conversion in liver of three patients and the effect of dietary methionine. Pediatr Res. 1970;4(3):238–238. 10.1203/00006450-197005000-00002

24. Pai YJ, Leung KY, Savery D, et al. Glycine decarboxylase deficiency causes neural tube defects and features of non-ketotic hyperglycinemia in mice. Nat Commun. 2015;6:6388. 10.1038/ncomms7388

25. Leung KY, Pai YJ, Chen Q, et al. Partitioning of One-Carbon Units in Folate and Methionine Metabolism Is Essential for Neural Tube Closure. Cell Rep. 2017;21(7):1795–1808. 10.1016/j.celrep.2017.10.072

26. Wolff JA, Kulovich S, Yu AL, Qiao CN, Nyhan WL. The effectiveness of benzoate in the management of seizures in nonketotic hyperglycinemia. Am J Dis Child. 1986;140(6):596–602. 10.1001/archpedi.1986.02140200106038

27. Van Hove JL, Vande Kerckhove K, Hennermann JB, et al. Benzoate treatment and the glycine index in nonketotic hyperglycinaemia. J Inherit Metab Dis. 2005;28(5):651–651. 10.1007/s10545-005-0033-x

28. Shelkowitz E, Saneto RP, Al-Hertani W, et al. Ketogenic diet as a glycine lowering therapy in nonketotic hyperglycinemia and impact on brain glycine levels. Orphanet J Rare Dis. 2022;17(1):423. 10.1186/s13023-022-02581-6

29. Hamosh A, McDonald JW, Valle D, Francomano CA, Niedermeyer E, Johnston MV. Dextromethorphan and high-dose benzoate therapy for nonketotic hyperglycinemia in an infant. J Pediatr. 1992;121(1):131–131. 10.1016/s0022-3476(05)82559-4

30. Hamosh A, Maher JF, Bellus GA, Rasmussen SA, Johnston MV. Long-term use of high-dose benzoate and dextromethorphan for the treatment of nonketotic hyperglycinemia. J Pediatr. 1998;132(4):709–709. 10.1016/s0022-3476(98)70365-8

31. Bjoraker KJ, Swanson MA, Coughlin CR, 2nd, et al. Neurodevelopmental Outcome and Treatment Efficacy of Benzoate and Dextromethorphan in Siblings with Attenuated Nonketotic Hyperglycinemia. J Pediatr. 2016;170:234–239. 10.1016/j.jpeds.2015.12.027

32. Korman SH, Wexler ID, Gutman A, Rolland MO, Kanno J, Kure S. Treatment from birth of nonketotic hyperglycinemia due to a novel GLDC mutation. Ann Neurol. 2006;59(2):411–415. 10.1002/ana.20759

33. Zammarchi E, Donati MA, Ciani F, Pasquini E, Pela I, Fiorini P. Failure of early dextromethorphan and sodium benzoate therapy in an infant with nonketotic hyperglycinemia. Neuropediatrics. 1994;25(5):274–274. 10.1055/s-2008-1073037

34. Boneh A, Allan S, Mendelson D, Spriggs M, Gillam LH, Korman SH. Clinical, ethical and legal considerations in the treatment of newborns with non-ketotic hyperglycinaemia. Mol Genet Metab. 2008;94(2):143–143. 10.1016/j.ymgme.2008.02.010

35. Stence NV, Fenton LZ, Levek C, et al. Brain imaging in classic nonketotic hyperglycinemia: Quantitative analysis and relation to phenotype. J Inherit Metab Dis. 2019;42(3):438–438. 10.1002/jimd.12072

36. Van Hove JL, Kishnani PS, Demaerel P, et al. Acute hydrocephalus in nonketotic hyperglycinemia. Neurology. 2000;54(3):754–754. 10.1212/wnl.54.3.754

37. Ichinohe A, Kojima K, Aoki Y, et al. Seizures and brain malformations in model mice for glycine encephalopathy. International Congress of Inborn Errors of Metabolism, September 2013; 2013; Brisbane, Australia.

38. Oda M, Kure S, Sugawara T, et al. Direct correlation between ischemic injury and extracellular glycine concentration in mice with genetically altered activities of the glycine cleavage multienzyme system. Stroke. 2007;38(7):2157–2157. 10.1161/STROKEAHA.106.477026

39. Kojima-ishii K, Kure S, Ichinohe A, et al. Model mice for mild-form glycine encephalopathy: behavioral and biochemical characterizations and efficacy of antagonists for the glycine binding site of N-methyl D-aspartate receptor. Pediatr Res. 2008;64(3):228–228. 10.1203/PDR.0b013e3181799562

40. Narisawa A, Komatsuzaki S, Kikuchi A, et al. Mutations in genes encoding the glycine cleavage system predispose to neural tube defects in mice and humans. Hum Mol Genet. 2012;21(7):1496–1496. 10.1093/hmg/ddr585

41. Santos C, Pai YJ, Mahmood MR, et al. Impaired folate 1-carbon metabolism causes formate-preventable hydrocephalus in glycine decarboxylase-deficient mice. J Clin Invest. 2020;130(3):1446–1446. 10.1172/JCI132360

42. Farris J, Alam MS, Rajashekara AM, Haldar K. Genomic analyses of glycine decarboxylase neurogenic mutations yield a large-scale prediction model for prenatal disease. PLoS Genet. 2021;17(2):e1009307. 10.1371/journal.pgen.1009307

43. Trask H, Tomlinson C, Fiering S, Gorham J, Muirhead K. Introducing DartMouse: the mouse speed congenic facility at Dartmouth Medical School. J Biomol Tech. 2010;21(3 Suppl):S70.

44. Wong GT. Speed congenics: applications for transgenic and knock-out mouse strains. Neuropeptides. 2002;36(2-3):230–236. 10.1054/npep.2002.0905

45. Toone JR, Applegarth DA, Levy HL. Prenatal diagnosis of non-ketotic hyperglycinaemia: experience in 50 at-risk pregnancies. J Inherit Metab Dis. 1994;17(3):342–342. 10.1007/BF00711825

46. Swanson MA, Miller K, Young SP, et al. Cerebrospinal fluid amino acids glycine, serine, and threonine in nonketotic hyperglycinemia. J Inherit Metab Dis. 2022;45(4):734–734. 10.1002/jimd.12500

47. Grant SL, Shulman Y, Tibbo P, Hampson DR, Baker GB. Determination of d-serine and related neuroactive amino acids in human plasma by high-performance liquid chromatography with fluorimetric detection. J Chromatogr B Analyt Technol Biomed Life Sci. 2006;844(2):278–278. 10.1016/j.jchromb.2006.07.022

48. Grabenstatter HL, Del Angel YC, Carlsen J, et al. The effect of STAT3 inhibition on status epilepticus and subsequent spontaneous seizures in the pilocarpine model of acquired epilepsy. Neurobiol Dis. 2014;62:73–85. 10.1016/j.nbd.2013.09.003

49. Grabenstatter HL, Carlsen J, Raol YH, et al. Acute administration of the small-molecule p75(NTR) ligand does not prevent hippocampal neuron loss or development of spontaneous seizures after pilocarpine-induced status epilepticus. J Neurosci Res. 2014;92(10):1307–1307. 10.1002/jnr.23402

50. Raible DJ, Frey LC, Del Angel YC, et al. JAK/STAT pathway regulation of GABAA receptor expression after differing severities of experimental TBI. Exp Neurol. 2015;271:445–456. 10.1016/j.expneurol.2015.07.001

51. Kane O, McCoy A, Jada R, et al. Characterization of spontaneous seizures and EEG abnormalities in a mouse model of the human A350V IQSEC2 mutation and identification of a possible target for precision medicine based therapy. Epilepsy Res. 2022;182:106907. 10.1016/j.eplepsyres.2022.106907

52. Mulcahey PJ, Tang S, Takano H, et al. Aged heterozygous Cdkl5 mutant mice exhibit spontaneous epileptic spasms. Exp Neurol. 2020;332:113388. 10.1016/j.expneurol.2020.113388

53. Dillon GM, Qu X, Marcus JN, Dodart J-C. Excitotoxic lesions restricted to the dorsal CA1 field of the hippocampus impair spatial memory and extinction learning in C57BL/6 mice. Neurobiology of Learning and Memory. 2008;90(2):426–426. 10.1016/j.nlm.2008.05.008

54. Mesches MH, Gemma C, Veng LM, et al. Sulindac improves memory and increases NMDA receptor subunits in aged Fischer 344 rats. Neurobiol Aging. 2004;25(3):315–315. 10.1016/S0197-4580(03)00116-7

55. Nielsen DM, Evans JJ, Derber WJ, et al. Mouse model of fragile X syndrome: behavioral and hormonal response to stressors. Behav Neurosci. 2009;123(3):677–677. 10.1037/a0015242

56. Cornejo BJ, Mesches MH, Benke TA. A single early-life seizure impairs short-term memory but does not alter spatial learning, recognition memory, or anxiety. Epilepsy Behav. 2008;13(4):585–585. 10.1016/j.yebeh.2008.07.002

57. Burman MA, Simmons CA, Hughes M, Lei L. Developing and validating trace fear conditioning protocols in C57BL/6 mice. J Neurosci Methods. 2014;222:111–117. 10.1016/j.jneumeth.2013.11.005

58. Hyde LA, Hoplight BJ, Denenberg VH. Water version of the radial-arm maze: learning in three inbred strains of mice. Brain Res. 1998;785(2):236–236. 10.1016/s0006-8993(97)01417-0

59. DeVore GR. Computing the Z Score and Centiles for Cross-sectional Analysis: A Practical Approach. J Ultrasound Med. 2017;36(3):459–459. 10.7863/ultra.16.03025

60. Autuori MC, Pai YJ, Stuckey DJ, et al. Use of high-frequency ultrasound to study the prenatal development of cranial neural tube defects and hydrocephalus in Gldc-deficient mice. Prenat Diagn. 2017;37(3):273–273. 10.1002/pd.5004

61. Walsh RN, Cummins RA. The Open-Field Test: a critical review. Psychol Bull. 1976;83(3):482–482.

62. Löscher W, Ferland RJ, Ferraro TN. The relevance of inter-and intrastrain differences in mice and rats and their implications for models of seizures and epilepsy. Epilepsy Behav. 2017;73:214–235. 10.1016/j.yebeh.2017.05.040

63. Brooks SP, Pask T, Jones L, Dunnett SB. Behavioural profiles of inbred mouse strains used as transgenic backgrounds. II: cognitive tests. Genes Brain Behav. 2005;4(5):307–317. 10.1111/j.1601-183X.2004.00109.x

64. Brooks SP, Pask T, Jones L, Dunnett SB. Behavioural profiles of inbred mouse strains used as transgenic backgrounds. I: motor tests. Genes Brain Behav. 2004;3(4):206–206. 10.1111/j.1601-183X.2004.00072.x

65. Bachmann C, Mihatsch MJ, Baumgartner RE, et al. [Non-ketotic hyperglycinemia: peracute course in neonatal period]. Helv Paediatr Acta. 1971;26(3):228–228.

66. Vogel P, Read RW, Hansen GM, et al. Congenital hydrocephalus in genetically engineered mice. Vet Pathol. 2012;49(1):166–166. 10.1177/0300985811415708

67. Jax Notes. Hydrocephalus in laboratory mice: The Jackson Laboratory; 2003 [updated 07/12/2003]. Available from: https://www.jax.org/news-and-insights/2003/july/hydrocephalus-in-laboratory-mice

68. Hoover-Fong JE, Shah S, Van Hove JL, Applegarth D, Toone J, Hamosh A. Natural history of nonketotic hyperglycinemia in 65 patients. Neurology. 2004;63(10):1847–1847. 10.1212/01.wnl.0000144270.83080.29

69. Tsuyusaki Y, Shimbo H, Wada T, et al. Paradoxical increase in seizure frequency with valproate in nonketotic hyperglycinemia. Brain Dev. 2012;34(1):72–72. 10.1016/j.braindev.2011.01.005

70. Yis U, Kurul SH, Dirik E. Nonketotic hyperglycinemia and acquired hydrocephalus. Pediatr Neurol. 2009;40(2):138–138. 10.1016/j.pediatrneurol.2008.10.007

71. Momb J, Appling DR. Mitochondrial one-carbon metabolism and neural tube defects. Birth Defects Res A Clin Mol Teratol. 2014;100(8):576–576. 10.1002/bdra.23268

72. Boneh A, Degani Y, Harari M. Prognostic clues and outcome of early treatment of nonketotic hyperglycinemia. Pediatr Neurol. 1996;15(2):137–137. 10.1016/0887-8994(96)00158-0

73. Festing M. Inbred Strains of Mice: 129 Bar Harbor: The Jackson Laboratory; 1976 [updated 04/09/1998]. Available from: http://www.informatics.jax.org/inbred_strains/mouse/docs/129.shtml.

74. Stein MR. The new generation of liquid intravenous immunoglobulin formulations in patient care: a comparison of intravenous immunoglobulins. Postgrad Med. 2010;122(5):176–176. 10.3810/pgm.2010.09.2214

75. Van Hove JL, Kishnani P, Muenzer J, et al. Benzoate therapy and carnitine deficiency in non-ketotic hyperglycinemia. Am J Med Genet. 1995;59(4):444–444. 10.1002/ajmg.1320590410

76. Kuseyri Hubschmann O, Julia-Palacios NA, Olivella M, et al. Integrative Approach to Predict Severity in Nonketotic Hyperglycinemia. Ann Neurol. 2022;92(2):292–292. 10.1002/ana.26423

77. Duran-Trio L, Fernandes-Pires G, Grosse J, et al. Creatine transporter-deficient rat model shows motor dysfunction, cerebellar alterations, and muscle creatine deficiency without muscle atrophy. J Inherit Metab Dis. 2022;45(2):278–278. 10.1002/jimd.12470

78. Meek S, Thomson AJ, Sutherland L, et al. Reduced levels of dopamine and altered metabolism in brains of HPRT knock-out rats: a new rodent model of Lesch-Nyhan Disease. Sci Rep. 2016;6:25592. 10.1038/srep25592

79. Rasmussen SA, Daenzer JMI, MacWilliams JA, et al. A galactose-1-phosphate uridylyltransferase-null rat model of classic galactosemia mimics relevant patient outcomes and reveals tissue-specific and longitudinal differences in galactose metabolism. J Inherit Metab Dis. 2020;43(3):518–518. 10.1002/jimd.12205

## Supplemental References

1. Trask H, Tomlinson C, Fiering S, Gorham J, Muirhead K. Introducing DartMouse: the mouse speed congenic facility at Dartmouth Medical School. J Biomol Tech. 2010;21(3 Suppl):S70.

2. Swanson MA, Coughlin CR, Jr., Scharer GH, et al. Biochemical and molecular predictors for prognosis in nonketotic hyperglycinemia. Ann Neurol. 2015;78(4):606–606. 10.1002/ana.24485

3. Toone JR, Applegarth DA, Levy HL. Prenatal diagnosis of non-ketotic hyperglycinaemia: experience in 50 at-risk pregnancies. J Inherit Metab Dis. 1994;17(3):342–342. 10.1007/BF00711825

4. Swanson MA, Miller K, Young SP, et al. Cerebrospinal fluid amino acids glycine, serine, and threonine in nonketotic hyperglycinemia. J Inherit Metab Dis. 2022;45(4):734–734. 10.1002/jimd.12500

5. Grant SL, Shulman Y, Tibbo P, Hampson DR, Baker GB. Determination of d-serine and related neuroactive amino acids in human plasma by high-performance liquid chromatography with fluorimetric detection. J Chromatogr B Analyt Technol Biomed Life Sci. 2006;844(2):278–278. 10.1016/j.jchromb.2006.07.022

6. Grabenstatter HL, Del Angel YC, Carlsen J, et al. The effect of STAT3 inhibition on status epilepticus and subsequent spontaneous seizures in the pilocarpine model of acquired epilepsy. Neurobiol Dis. 2014;62:73–85. 10.1016/j.nbd.2013.09.003

7. Grabenstatter HL, Carlsen J, Raol YH, et al. Acute administration of the small-molecule p75(NTR) ligand does not prevent hippocampal neuron loss or development of spontaneous seizures after pilocarpine-induced status epilepticus. J Neurosci Res. 2014;92(10):1307–1307. 10.1002/jnr.23402

8. Raible DJ, Frey LC, Del Angel YC, et al. JAK/STAT pathway regulation of GABAA receptor expression after differing severities of experimental TBI. Exp Neurol. 2015;271:445–456. 10.1016/j.expneurol.2015.07.001

9. Kane O, McCoy A, Jada R, et al. Characterization of spontaneous seizures and EEG abnormalities in a mouse model of the human A350V IQSEC2 mutation and identification of a possible target for precision medicine based therapy. Epilepsy Res. 2022;182:106907. 10.1016/j.eplepsyres.2022.106907

10. Mulcahey PJ, Tang S, Takano H, et al. Aged heterozygous Cdkl5 mutant mice exhibit spontaneous epileptic spasms. Exp Neurol. 2020;332:113388. 10.1016/j.expneurol.2020.113388

11. Walsh RN, Cummins RA. The Open-Field Test: a critical review. Psychol Bull. 1976;83(3):482–504.

12. Kojima-ishii K, Kure S, Ichinohe A, et al. Model mice for mild-form glycine encephalopathy: behavioral and biochemical characterizations and efficacy of antagonists for the glycine binding site of N-methyl D-aspartate receptor. Pediatr Res. 2008;64(3):228–228. 10.1203/PDR.0b013e3181799562

13. Mesches MH, Gemma C, Veng LM, et al. Sulindac improves memory and increases NMDA receptor subunits in aged Fischer 344 rats. Neurobiol Aging. 2004;25(3):315–315. 10.1016/S0197-4580(03)00116-7

14. Nielsen DM, Evans JJ, Derber WJ, et al. Mouse model of fragile X syndrome: behavioral and hormonal response to stressors. Behav Neurosci. 2009;123(3):677–677. 10.1037/a0015242

15. Cornejo BJ, Mesches MH, Benke TA. A single early-life seizure impairs short-term memory but does not alter spatial learning, recognition memory, or anxiety. Epilepsy Behav. 2008;13(4):585–592. 10.1016/j.yebeh.2008.07.002

16. Burman MA, Simmons CA, Hughes M, Lei L. Developing and validating trace fear conditioning protocols in C57BL/6 mice. J Neurosci Methods. 2014;222:111–117. 10.1016/j.jneumeth.2013.11.005

17. Hyde LA, Hoplight BJ, Denenberg VH. Water version of the radial-arm maze: learning in three inbred strains of mice. Brain Res. 1998;785(2):236–236. 10.1016/s0006-8993(97)01417-0

